# The last days of *Aporia crataegi* (L.) in Britain: evaluating genomic erosion in an extirpated butterfly

**DOI:** 10.1101/2023.12.19.572305

**Authors:** Rebecca Whitla, Korneel Hens, James Hogan, Geoff Martin, Casper Breuker, Timothy G. Shreeve, Saad Arif

## Abstract

Current rates of habitat degradation and climate change are causing unprecedented declines in global biodiversity. Studies on vertebrates highlight how conservation genomics can be an effective tool to identify and manage threatened populations, but it is unclear how vertebrate derived metrics of genomic erosion translate to invertebrate species, with their markedly different population sizes and life histories. The Black-veined White butterfly (*Aporia crataegi)* was extirpated from Britain in the 1920s. Here, we sequenced historical DNA from 17 museum specimens collected between 1854-1924 to reconstruct demography and compare levels of genomic erosion between extirpated British and extant European mainland populations. We contrast these results using modern samples of the Common Blue butterfly (*Polyommatus icarus*); a species with relatively stable demographic trends in Great Britain. We provide evidence for bottlenecks in both these species around the period of post-glacial colonisation of the British Isles. Our results reveal different demographic histories and *N_e_* for both species, consistent with their fates in Britain, likely driven by differences in life history, ecology, genome lengths and body size. Despite a difference, by an order of magnitude, in historical effective population sizes (*N_e_*), reduction in genome-wide heterozygosity in *A. crataegi* was comparable to that in *P. icarus*. Symptomatic of *A. crataegi*’s disappearance were marked increases in runs-of-homozygosity (RoH), potentially indicative of recent inbreeding, and accumulation of putatively mildly and weakly deleterious variants. Our results support the idea that metrics of inbreeding and accumulation of deleterious mutations could be more informative markers of population or species in decline.

## Introduction

Global extinction rates have increased over the last 500 years (Barnosky *et al*., 2011, Ceballos *et al*., 2015, Cowie *et al*., 2022), accelerating over the last 100 years with *circa* 25% of species recently assessed by the IUCN are now threatened with extinction (IPBES, 2019). In Great Britain, *c.* 16.1% of assessed flora and fauna species (10,008) are classified as threatened, and 146 species have gone extinct (with 52 species ‘likely extinct’) since 1500 (Burns *et al*., 2023). Invertebrates count for 97% of all animals globally and are a major component of terrestrial biodiversity with important roles in ecosystem functioning. Assessments for many invertebrates are data deficient, precluding accurate determination of their status (Karam-Gemael *et al*., 2020). Reasons for insufficient data include underfunding for less charismatic and economically important groups (Cardoso and Leather, 2019) and difficulties in sampling and identification (Wagner *et al*., 2021).

Among insects, butterflies are one of the best-studied groups (for example, Warren, 1992; Thomas, 2005; Dincă *et al*., 2015; Middleton-Welling *et al*., 2020; Warren *et al*., 2021), can be used as bioindicators of environmental and climate change (Dennis *et al*., 2003; Hill *et al*., 2021) and serve as an important invertebrate model for communicating science to society (Preston *et al*., 2021). Since the 1970s, standardised and systematic monitoring of butterflies across Europe has provided a comprehensive and continuous record of changes in biodiversity. Compared to most of continental Europe, a larger proportion of UK butterflies are now either extirpated (4 of 62 species) or considered as critically endangered or vulnerable (19/62) (Fox *et al*., 2022).

By quantifying genomic erosion, conservation genomics can be a valuable tool in conservation, helping identify vulnerable populations and informing translocations and/or breeding programmes (Díez-del-Molino *et al.,* 2018; Bosse and van Loon, 2022). Metrics for genomic erosion, associated with reductions of fitness (Bosse and van Loon, 2022), aim to estimate genetic diversity, effective population size (*N_e_*), levels of inbreeding and genetic load. All have a strong theoretical basis to be correlated with and potentially exacerbate extinction risk in small and vulnerable populations (Gomulkiewicz and Holt, 1995; Lynch *et al*., 1995a; Charlesworth and Willis, 2009). Negative impact of inbreeding depression in small populations is well established (Saccheri *et al*., 1998; Mattila *et al*., 2012; Neaves *et al*., 2015; Rowe *et al.,* 2017) but the utility of such metrics of, particularly those relying solely on genome-wide, putatively neutral genetic markers, in predicting extinction risk is currently under debate (Kardos *et al*., 2016; Teixeira and Huber, 2021; Dussex *et al*., 2023). A better understanding is emerging from the accumulation of genomic data of small and threatened populations using whole genomes to estimate multiple metrics of genomic erosion simultaneously (DeWoody *et al*., 2021; Bosse and van Loon, 2022; Dussex *et al*., 2023), but is largely limited to examples from terrestrial vertebrates and our understanding of genomic erosion in insects is far more limited (Webster *et al*., 2023; but see de-Dios *et al*., 2023; Bortoluzzi *et al*., 2023). Insects can have markedly higher population sizes and different life histories, compared to vertebrates, and it is unclear how this would impact readouts of genetic diversity, effective population size (*N_e_*), levels of inbreeding and genetic load in threatened insect populations. Comparative analyses of genomic erosion between endangered and stable or expanding insect populations, could prove extremely useful for detecting insect declines or identifying vulnerable species or populations from small or limited samples and for determining effective metrics to identify these in insects.

Here, we examine trends of genomic erosion in *Aporia crataegi* (L.) (Black-veined White butterfly), which was extirpated from the British Isles *c*. 1925 (Pratt, 1983), contrasting these with populations in continental Europe. We also provide a comparison between British and continental European populations of *Polyommatus icarus* (Rott.) (Common Blue butterfly); a species that has exhibited little change in long term population trends (https://www.gov.uk/government/statistics/butterflies-in-the-wider-countryside-uk). *A. crataegi* is widespread throughout much of Europe, Asia and north Africa. It has recently been lost from the Netherlands, Czechia (van Swaay *et al*., 2010), and South Korea (Kim *et al*., 2015), and is declining in other European countries (van Swaay *et al*., 2010). Prior to extinction in Britain, it was restricted to southern and central England and Wales and there is evidence that populations were in decline from the mid to late 19^th^ century (Dale, 1887; South 1906; Pratt, 1983) and was primarily only found in southeast England by the end of the 19^th^ Century. Its decline in Europe is typically attributed to habitat loss, but this does not seem to be the case for its extirpation in Britain (MacLachlan, 1893) with pesticide use, parasites, microbial or fungal pathogens and predation by small birds all being suggested as potential reasons instead (reviewed in Pratt, 1983). Numerous “accidental” releases, recorded as early as the 19^th^ century (Pratt, 1983), from butterfly enthusiasts to reintroduce the species from continental European stock have been made, but none have been successful for establishment of the species.

To examine whether *A. crataegi* in Britain exhibited any symptoms of genomic erosion, we sequenced historical specimens of *A. crataegi* from Britain and continental Europe dating from 1854-1924. We specifically addressed whether the British *A. crataegi* were distinct from continental Europe based on genetic structuring and demographic history prior to 10 Kya. We examine individual heterozygosity, runs-of-homozygosity ((RoH), a genomic measure of inbreeding), and genetic load to explain the vulnerability of the British Isles populations prior to extinction and contextualise these finding by comparison with corresponding analyses of the relatively stable British and continental European *P. icarus*.

## Materials and Methods

### Historical A. crataegi Specimen Selection

Seventeen *A. crataegi* specimens from Britain (GB), Belgium, and France, (Fig. 1A, Table S1) were sampled from collections at Oxford University Museum of Natural History (OUMNH), Oxford, England and the Natural History Museum (NHM), London, England. Specimen dates ranged from 1854-1924. For comparison, we only used museum samples of continental populations to avoid any potential bias stemming from comparing degraded historical DNA with modern DNA. Up to two legs per specimen were taken from pinned samples for DNA extraction, although in some cases only a single leg was available.

**Figure 1:**
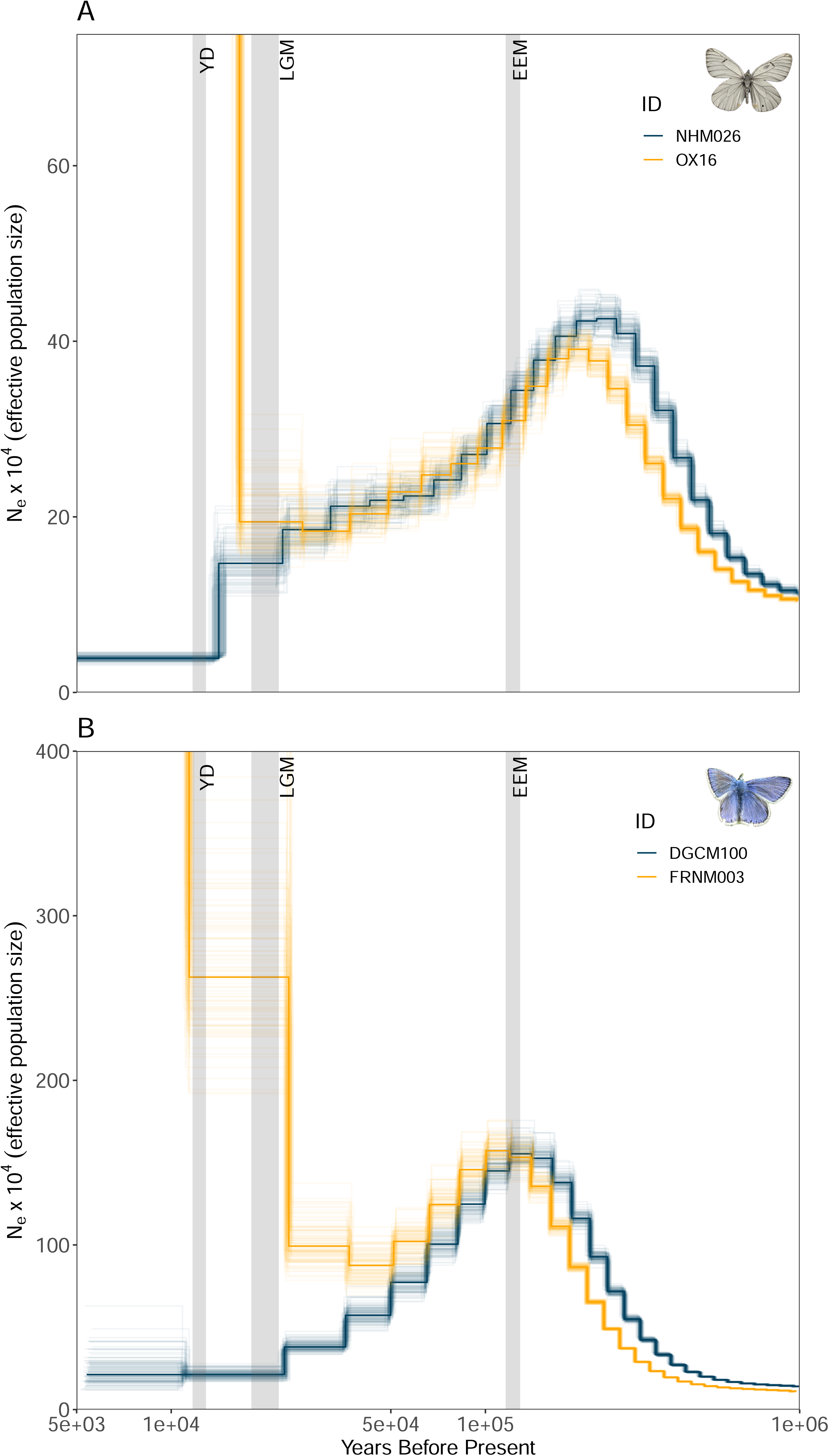
(A): Collection locations (based on museum records) of historical A. crataegi specimens (circles) and of modern P. icarus (triangles). NHM prefix refers to specimens obtained from Natural History Museum, London and OX for those obtained from Oxford University Natural History Museum, Oxford. (B): Principal component analysis (PCA) for A. crataegi SNPs. The PCA is based on 14,930 LD-pruned SNPs based on hard genotype calls. A PCA using genotype likelihoods shows a similar pattern (Fig S3). Asterisks indicate UK-collected specimens that have clustered with European specimens. (C): PCA for P. icarus specimens based on 197,182 LD-pruned SNPs showing similar differentiation between British and European specimens. Inset images: A pinned A. crataegi specimen and live P. icarus in their respective plots.

### Extraction, Library Preparation and Sequencing of Historical DNA

DNA extractions were undertaken on a laboratory bench used only for historical DNA (hDNA) extractions to prevent contamination from modern DNA. Extraction blanks were prepared alongside samples to monitor potential contamination. Legs of specimens were fully macerated using a Tissuelyser II bead mill (Qiagen) and extracted using a QIAamp DNA Micro kit (Qiagen) using the standard protocol with a final elution volume of 20μL. Total DNA yield was measured using a Qubit 4 Fluorometer (Life Technologies) with a Qubit dsDNA HS Assay Kit. Eluted DNA was stored in Eppendorf DNA LoBind Tubes at -20°C until library preparation.

Genomic library preparation of hDNA was performed using the NEBNext Ultra II DNA Library Prep Kit for Illumina. A CUT&RUN (Liu, 2019) protocol was used, modified for use with low yield samples and samples with short fragment sizes. NEBNext® Dual Index Multiplex Oligos for Illumina (Set 1) Primers were used to construct multiplexed libraries. SPRIselect beads (Beckman Coulter) were used for size exclusion post primer ligation at 1.75X or 2.2X concentration.

The number of PCR cycles used for amplification was dependent on the input DNA quantity; 12 cycles for samples < 5 ng and 10 cycles for samples ε 5 ng. Libraries were then purified with 1.2X SPRIselect beads to remove adapter dimer. Library enrichment was performed using either a Q5U® Hot Start High-Fidelity DNA Polymerase (New England Biolabs) or the NEBNext Ultra II Q5DNA polymerase (Table S1). We did not attempt to repair the hDNA template prior to library preparation. However, the NEBNext Ultra II DNA Library Prep Kit makes use of USER enzyme during the adapter ligation step. This excises Uracil bases from the ligated adapters and is required when using NEBNext adapters, but in case of hDNA template with post-mortem deamination damage, the enzyme will also excise Uracil bases on the 5’ end of the hDNA template. With subsequent paired-end Illumina sequencing this results in partial repair with 5’ ends exhibiting little to no signs of historical damage whereas the damage profile on 3’ ends reflects typical post-mortem DNA damage.

Quality control for the libraries was performed using a Qubit 4 Fluorometer (Life Technologies) and an Agilent Bioanalyser, using the Agilent High Sensitivity DNA analysis Chip Kit. Sample libraries were pooled equimolarly, with extraction blanks pooled at a 1:10 molar ratio in proportion to sample libraries. Sequencing was performed using the NovaSeq Illumina platform using a 150 base pair (bp) paired-end (150PE) strategy at Novogene (Cambridge).

### Sampling Modern specimens of P. icarus

Four individuals of *P. icarus* were collected from across the British Isles and an additional three from Averyon, France between 2017-2018 (Table S2, Fig. 1A) and DNA extraction and further details of these samples is as described in Arif *et al*. (2021). DNA was normalised to 10 ng/ul and library preparation and sequencing, to a theoretical coverage of 15-50X (Table S2), was performed using the NovaSeq Illumina platform with a 150PE strategy at Novogene (Cambridge).

### Data Processing

The GenErode pipeline v0.4.2 (Kutschera *et al*., 2022), suitable for processing both modern and historical short-read data, was used to trim and process all modern and historical sequence data along with highly contiguous reference genomes for *A. crataegi* (Ebdon *et al*., 2022) or *P. icarus* (Lohse *et al*., 2023). Both these reference genome assemblies are high quality, produced from single male specimens, with an assembled Z chromosome (Lepidopteran males are commonly the homogametic sex). The *A. crataegi* assembly is ∼230 Megabases (Mb) long and is scaffolded into 26 pseudochromosomes and a Z chromosome. The *P. icarus* assembly is ∼512 Mb scaffolded into 23 pseudochromosomes and a Z.

FastQC v0.11.9 (Andrews *et al*., 2012) and fastp v0.20.1 (Chen *et al*., 2018) were used to trim adapters and merge paired sequences (for historical samples). Trimmed reads were aligned to reference genomes using BWA aln for hDNA and BWA mem for modern data (Li and Durbin, 2009) with default settings in the GenErode pipeline. Post-mapping, any samples sequenced across multiple lanes were merged and PCR duplicates removed. MapDamage v2.0 (Jónsson *et al*., 2013) was used to estimate damage in historical samples and rescale base quality scores. Next, GATK IndelRealigner, (Van der Auwera and O’Connor, 2020) was used to remap any reads around indels to improve mapping accuracy. For the modern, *P. icarus* data, all resulting BAM files were subsampled to 10X to mitigate the influence of varying coverage on heterozygous calls. These BAM files were processed further to call variants and reconstruct historical demography. For the latter, we used full coverage data for the modern *P. icarus* samples.

Variants calls were made on a per-sample basis using Bcftools mpileup (Danecek *et al*., 2021) with a minimum depth threshold of 1/3 of average coverage (with an absolute minimum of 3X) and a maximum of ten times the average coverage while minimum mapping quality and minimum base quality were both set to 30 (-Q 30 -q 30). Additional filters as a part of the GenErode pipeline, included an allelic imbalance filter for heterozygous sites, removal of variants in repetitive regions of the genome, and removal of variants in CpG regions. A final VCF was generated that consisted of only biallelic single nucleotide polymorphisms (SNPS) and genotypes with no more than 10% missing data across all samples (historical *A. crataegi* samples, containing 305,394 SNPs) or no missing data (modern *P .icarus* data, containing 5,656,704 SNPs). SNPs from these VCFs were used for downstream analyses, unless otherwise stated. Our configuration files for the GenErode pipeline are available at https://github.com/rmwhitla/BVWpaper.

### Population Structure

Population structure of both the historical *A. crataegi* and modern *P. icarus* samples were compared using principal component analysis (PCA) based on filtered genotypes in VCF format. Before PCA, any SNPs on the Z chromosome were filtered out. Furthermore, we pruned the remaining SNP sets for linkage disequilibrium over windows of 10 kilobases with a step size of 10 bases and an *r^2^* of 0.1 using Plink 1.9 (Purcell *et al.,* 2007) resulting in a total of 14,930 (*A. crataegi*) or 197,182 *(P. icarus*) SNPs. PCA was performed using Plink v1.9 with a minor allele frequency (MAF) threshold of 10% (historical *A. crataegi)* or 15% (modern *P. icarus*).

Given the low and variable coverage (2-13X) of the historical *A*. *crataegi* samples, we also performed PCA based on genotype likelihoods, which accounts for statistical uncertainty in called genotypes arising from low coverage data (Korneliussen *et al*., 2014). Genotype likelihoods were calculated using ANGSD (Korneliussen *et al*., 2014) from BAM files produced as part of the GenErode pipeline with a minimum mapping quality at 30 and minimum base quality at 20 (-minMapQ 30 -minQ 20) filtering only for the same sites used with the PCA for hard genotype calls. Pcangsd (Meisner and Albrechtsen, 2018) was used for the PCA of genotype likelihoods with a minimum allele frequency of 10% (--minMaf 0.1).

### Historical Demography Reconstruction

We used the Pairwise Sequentially Markovian Coalescent model as implemented in PSMC v0.6.5 (Li and Durbin, 2011) to determine if British populations of *A. crataegi* and *P. icarus* exhibit similar trends of historical change in effective population size (*N_e_*) following permanent colonization of the British Isles after the Younger Dryas *c.*11.5 Kya and permanent flooding of the Channel *c.* 9 Kya (Dennis, 1992). As input for PSMC, we generated a consensus fastq file from BAM files, using only samples with average coverage greater than 5X. Consensus fastq files were generated with Bcftools pileup and vcfutils.pl, filtering positions with minimum depth less than 5X or maximum depth greater than 100X in *A. crataegi* and with minimum depth less than 5X or maximum depth greater than 500X in *P. icarus*. Additionally, any positions in repeat masked regions or CpG regions (historical samples only) or Z chromosomes were also removed. PSMC was run with: number of iterations -N = 25, maximum coalescent time -t = 8 for *A. crataegi* or t = 9 fpr *P. icarus* and the atomic time interval (-p) at 66 (for *A. crataegi*: 4+25*2+3+9) and 66 (for *P. icarus*: 27*2+4+8). The atomic time interval and maximum coalescent time was chosen to avoid overfitting by ensuring that at least 10 recombination events occurred over each interval after the 20^th^ iteration of PSMC (Li and Durbin, 2011). To account for low coverage and stochastic loss of heterozygosity, the initial theta/rho ratio parameter for each sample was calculated based on false negative rates (FNR) corresponding to the coverage of that sample.

FNR for different coverages were determined based on the recommendation of Li and Durbin (2011) (https://github.com/lh3/psmc), and as implemented by Sarabia *et al*. (2020). To obtain FNR estimated for *A. crataegi*, we used a subsampled version of a publicly available high coverage (23X) modern *A. crataegi* (accession: SRR7948941) specimen from Japan that was sequentially subsampled to 5X, 8X, 10X, 15X and 20X. For FNR correction for low coverage *P. icarus* samples, we used the same subsampling strategy as for *A. crataegi* using one of our high coverage samples (DGCM100, coverage: >50X). The initial theta/rho (-r) parameter for PSMC was then set to 5/(1-FNR). To visualise the variability in our PSMC estimates, we ran 100 bootstraps for each sample over 5 Mb segments using the same initial parameters as the original run for each sample.

To scale population size and times resulting from the PSMC analysis, we used a direct estimate of spontaneous mutation rate of 2.9 x10^-9^ from *Heliconius melpomene* (Keightley *et al.,* 2015). For *A. crataegi*, which is univoltine in the geographic range our samples came from (Solonkin *et al.,* 2021), we used 1 generation per year. For *P. icarus*, we used 2 generations per year, although the number of broods are variable in this species (one in Scotland but two or three in Southern England and parts of Europe). We tested the sensitivity of the PSMC curves for *P. icarus* by changing the generation time to 1 and 3 broods per year.

### Genetic Diversity

Genome-wide heterozygosity was estimated using ANGSD, which utilises a genotype likelihood approach that is more suitable for low coverage data (Korneliussen *et al*., 2014). We calculated genotype likelihoods as described above (PCA using genotype likelihoods), except in this case no LD pruning was performed. To estimate global heterozygosity, we calculated the unfolded site frequency spectrum (SFS) using the realSFS function in ANGSD.

### Inbreeding and Contemporary *N_e_*

We estimated individual inbreeding coefficients based on the detection of the number and size of Runs of Homozygosity (RoH). RoH were calculated using Plink v1.9 (Purcell *et al.,* 2007). We used a sliding window size of 100 SNPs (homozyg-window-snp 100). The maximum number of heterozygotes per window was 2 (homozyg-window-het 2). Minimum SNP count for each RoH was 25 (homozyg-snp 25) and the minimum size was set to 50kb (homozyg-kb 50). We allowed 10 missing sites per window (intermediate filtering - homozyg-window-missing 10). For the maximum number of heterozygote sites within a RoH, we allowed 3 (homozyg-het3). Using these results, F_RoH_, the inbreeding coefficient, was determined using the overall proportion of the genome in RoH. Results from Plink were summarised in R v.4.3.0 (R Core Team, 2023) using the package DetectRuns (Biscarini *et al.,* 2018). F_ROH_ was estimated for RoH ε 100 Kilobases (Kb), RoH ε 500 Kb or RoH ε 1 Megabase (Mb) for samples with average coverage ε 5X. RoH ε 100 Kb and less than 1 Mb may result from background inbreeding due to drift whereas RoH ε 1 Mb result from recent consanguineous mating (Ceballos *et al*., 2018).

Inference of *N_e_* from PSMC is limited over the last 10 Kya but the length and distributions of RoHs can be informative of more recent *N_e_*. To compare our historical estimates of *N_e_* (∼10 Kya) estimated from PSMC with those observed from RoH we calculated the probability of observing at least one RoH ═ 1 Mb, in a genome of size 230 Mb (*A. crataegi*) or 512 Mb (*P. icarus).* Assuming a neutrally evolving Wright-Fisher population, the probability of a randomly chosen region of length 10^6^ bases being a ROH can be computed as the probability that pairwise coalescence happens before either sequence recombines (Hudson, 1990):

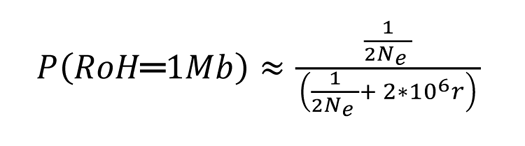

Where *N_e_* was based on the PSMC estimate at ∼10 Kya and *r* represents the recombination rate and was set to 20 times 2.9 x10^-9^ (the spontaneous mutation rate). This choice of recombination rate is empirically plausible for butterflies which can be 20-fold higher than the mutation rate. (e.g. Jiggins *et al*., 2005; Palahí i Torres *et al.,* 2023)

### Genetic Load Estimation

To estimate the number of deleterious mutations or the genetic load we used snpEff (Cingolani *et al.,* 2012) to annotate genetic variants of high impact (likely highly disruptive to protein, e.g., loss of function (LoF) variants), moderate impact (e.g. missense variants) and low impact (synonymous, likely harmless but may be weakly deleterious under strong codon bias), as well as modifier (non-coding) variants. We focused on annotated variants within coding regions (LoF, missense or synonymous). snpEFF only predicts the potential phenotypic impact of the variants, and we lack any prior knowledge of the selection or dominance coefficients for these variants.

For the analysis of genetic load, we only used samples with ε 5X average depth and no missing data across samples. This resulted in 179,135 SNPs for historical *A. crataegi* and 5,656,704 SNPs for *P. icarus*. Gene annotations (as GTF files) for both species were retrieved from Ensembl’s Darwin Tree of Life data portal (https://projects.ensembl.org/darwin-tree-of-life/). Both annotations were generated by the Ensembl Gene Annotation System (Aken *et al*., 2017) optimised for non-vertebrates. The annotation for *A. crataegi* (Ebdon *et al.,* 2022) is based on a single male from Catalunya, Spain and the annotation for *P. icarus* (Lohse, 2023) is based on a single male from Scotland.

We also determined the relative excess in the frequency of shared derived alleles (ignoring the impact of *de novo* mutations) across the LoF, missense and synonymous categories in both species. We used an approach previously described by Xue *et al*. (2015) as implemented by van der Valk *et al*. (2019). For each variant category we calculate a ratio *R_xy_* that is a measure of shared derived alleles with a higher frequency in population X (British) compared to population Y (European). An *R_xy_* > 1 implies a relative increase in frequency in population and X compared to Y, whereas < 1 implies a relative decrease in X relative to Y. To estimate confidence intervals for our *R_xy_* ratios we used a jackknife approach dropping a single chromosome at a time.

### Statistical Analysis

For all statistical analysis between historical specimens of *A. crataegi* we used a linear model with average coverage as a covariate to account for variation in coverage between specimens. Marginal means of all estimates for each group were estimated using the R package emmeans (Lenth, 2023). For *P. icarus*, statistical comparisons were made using Welch’s two sample t-tests. All statistical comparisons were performed in R v.4.3.0 (R Core Team, 2023).

## Results

### Resequencing of hDNA

The final sequencing depth of the 17 historical *A. crataegi* specimens varied between 2X-13X, with higher mean coverage of European samples (10.13X) compared to British specimens (6.03X) and sequencing depth varied from 2X-13X (Table S1). The average proportion of the genome sequenced to a depth of 5X or greater was 50.77% (Table S1).

Post-mortem degradation was apparent in the 3’ ends of our historical reads (Fig. S1) as an increase in G>A transversions. The 5’ ends of our fragments show little to no sign of historical damage due to the application of the USER enzyme during library preparation. Mean fragment length of sequencing reads, after trimming and merging reads, was 49bp with average fragment length per sample ranging from 44-65bp (Table S1). There was a significant positive correlation between mean fragment length and age of sample (Fig. S2) with fragment lengths increasing closer to more recent times (Mullin *et al.,* 2023).

In our extraction blanks (BL1 and BL2), 0.49% and 0.43% of reads mapped to the *A. crataegi* genome respectively. In the blanks, 0.2% of bases had 1X coverage, and 0% of bases had higher coverage. The average read lengths in the two blanks were 91 and 97. Overall, analysis of the mapping profile of the blanks suggests no evidence for cross contamination during DNA extraction and library preparation of the historical samples.

### Spatial Genetic Structuring between Great British and European Mainland Butterflies

A PCA using LD-pruned SNPs of all historical *A. crataegi* specimens identifies spatial genetic structuring across historical samples regardless of collection date (Fig. 1B). A PCA based on genotype likelihoods (Fig. S3), undertaken because of low to moderate coverage of the historic specimens, yielded visually similar results to the one based on hard genotype calls (Fig. 1B).

PC1 explained 10.8% of the variation, while PC2 explained 8.6% of variation. Spatial genetic structuring is apparent along PC1 with European samples tightly clustering on one end with negative values for PC1, whereas points with greater values of PC1 represent two specimens from range margins (Wales; see Fig. S3 for a labelled PCA- but note the axes are reversed) but also include a sample with unknown origins (K05L) and one specimen from Kent (NHM026). In contrast to the geographical cline apparent along PC1, there is no clear cline present along PC2 but the points with the most extreme values represent the most recent specimens from Kent (NHM928 and NHM019). *P. icarus* exhibit similar structuring along PC1 with the southern England sample (Fig. 1C) sorting in between the European and marginal Scottish samples. Overall, sorting of specimens along PC1 for both species reveals a tight clustering among European butterflies and more geographic structuring within GB samples.

Three historical *A. crataegi* specimens (NHM898, NHM027 and NHM918; Fig. 1B) labelled as GB clustered closer to French and Belgium samples rather than the GB samples. Rather than a case of mislabelling, these specimens may be potential European migrants or European specimens released in Britain. The latter was not an uncommon practise from the late 19^th^ to early 20^th^ century (Pratt, 1983). For subsequent analyses, these three samples are grouped with the other mainland European samples.

### Demographic reconstruction

Historical demographic reconstruction of *A. crataegi* using PSMC shows similar trajectories of effective population size (*N_e_;* Fig. 2A, Fig. S4) for both British and European samples up to the last glacial maximum (LGM, *c.* 22 to 17 Kya). British *A. crataegi,* show a marked decrease in *N_e_* around the LGM and Younger Dryas (12.8–11.5 Kya). A similar pattern of trajectories is also observed in *P. icarus* (Fig. 2B, Fig. S5) but, the divergence in *N_e_* appears to happen earlier, just before the LGM, but this trend is sensitive to the number of generations per year (Fig. S6). Both reconstructions indicate British *A. crataegi* and *P. icarus* show reduced *N_e_* before or around 12 Kya.

**Figure 2:**
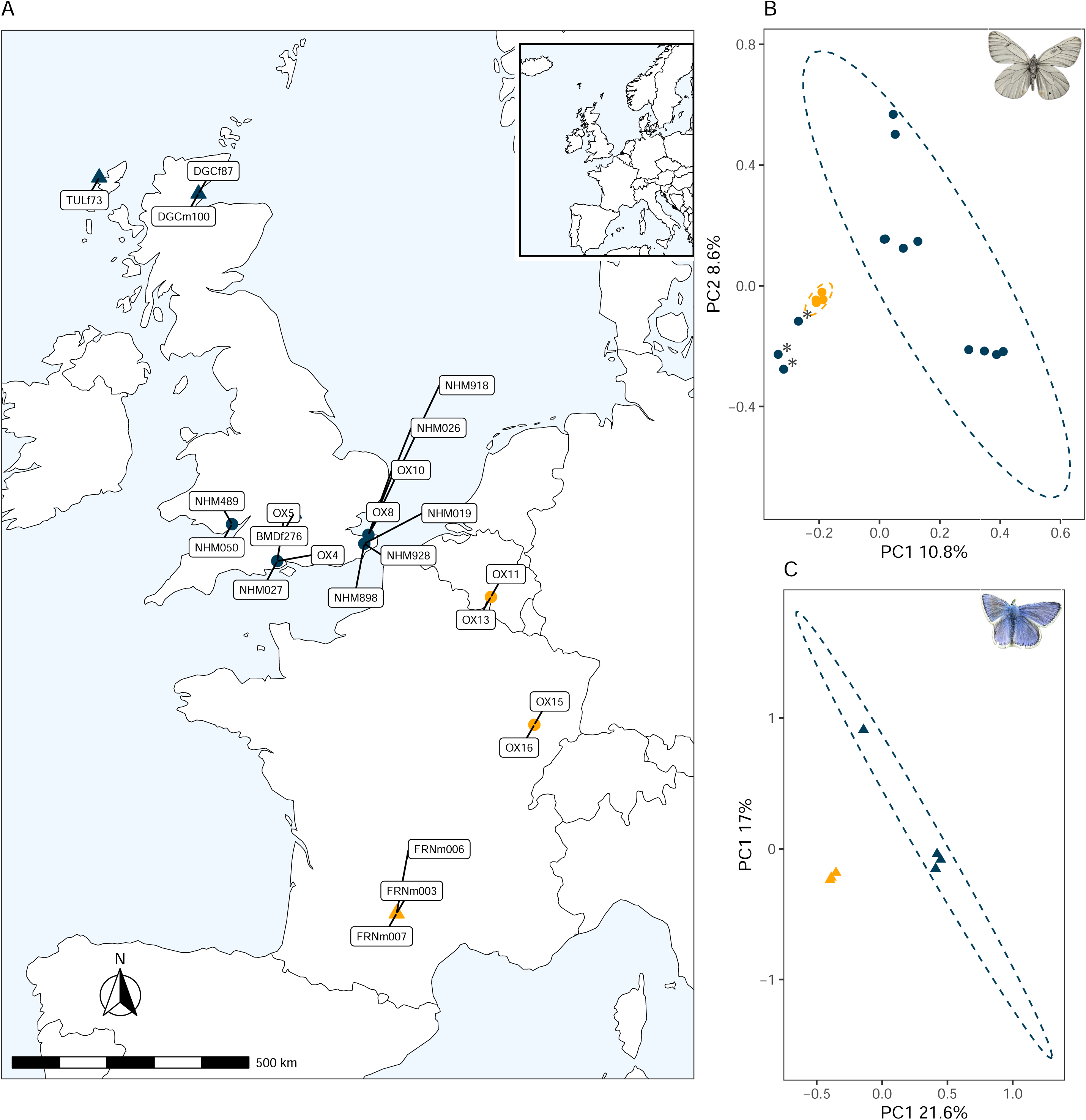
(A): PSMC plot for A. crataegi specimens, showing changing Ne over time and population divergence between a GB specimen (blue curve) and a European specimen (yellow curve). All A. crataegi specimens PSMC curves are shown in S4. (B): PSMC for modern P. icarus specimens, with a GB specimen (blue curve) and a European specimen (yellow curve). Divergence timing is shown as before the LGM due to interval settings during the PSMC run (-p). YD = Younger Dryas, LGM= Last Glacial Maximum, EEM= Eemian Interglacial Period

Approximation of *N_e_* at the time of the Younger Dryas (∼10 Kya) are 3.87 x 10^4^ for *A. crataegi,* and 2.14 x 10^5^ for *P. icarus* (Fig. 2). We note that although we used higher coverage samples to estimate the FNR for low coverage samples, for samples below 8X (*A. crataegi*) or 14X (*P. icarus*) we could not recover *N_e_* estimated for the high coverage data before 50 Kya (Fig. S7 and S8). However, this discrepancy was limited to the magnitude of *N_e_* and not the trajectories themselves.

### Estimates of heterozygosity

Individual genome-wide heterozygosity (Fig. 3A-B) rates were lower for British samples for both *A. crataegi* and *P. icarus*. Heterozygosity between British and European *A. crataegi* was significantly different (*F_1,14_*=25.21, *p*=0.000187, Fig. 3A), after accounting for variability in sample coverage. British *A. crataegi* exhibited an average reduction of 17.9% in heterozygosity relative to European samples. Heterozygosity was also significantly different between the British and French *P. icarus* (*t=*9.26, *degrees of freedom* (*df*) = 3.52, *p*= 0.001356; Fig. 3B) samples, with an average reduction of 14.5%.

**Figure 3:**
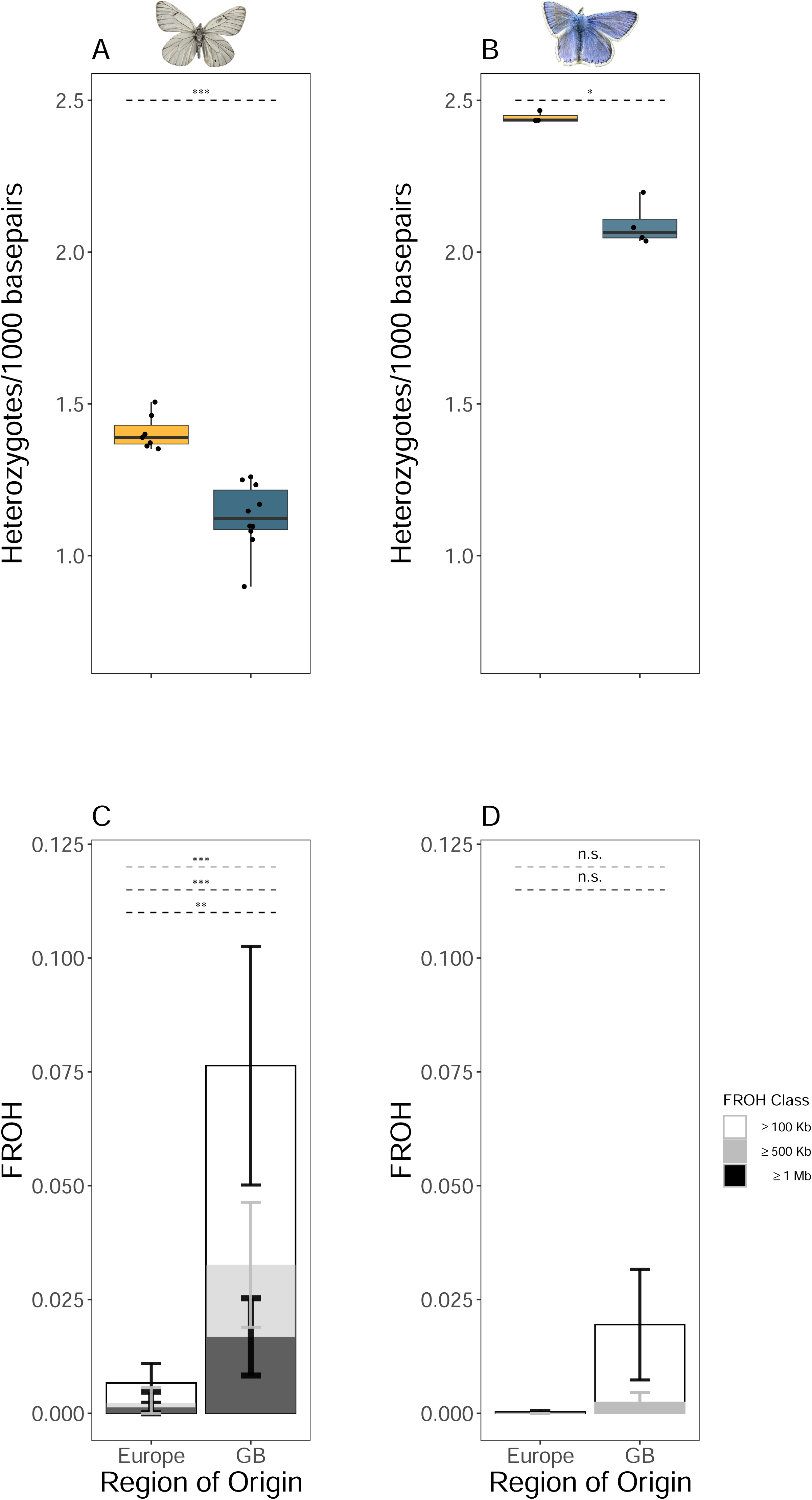
(A): Individual heterozygosity rates in A. crataegi specimens from Europe and GB, calculated using ANGSD. Dashed lines show heterozygosity difference between Europe and GB. Significance codes: n.s. = not significant, * = p <0.05, ** = p <0.01, *** = p < 0.001. (B): Individual heterozygosity rates in P.icarus specimens from Europe and GB (C): RoH in A. crataegi specimens from Europe and GB. Open bars show the total proportion of the genome in RoH ≥ 100 KB, light grey bars show RoH ≥ 500kb and dark grey bars show proportions in RoH ≥ 1 MB. (D): RoH in P.icarus European and GB specimens. Dashed lines in C and D show significance of difference between European and GB specimens.

### Inbreeding and concordance of historical and recent *N_e_*

For detecting RoHs we used three thresholds: ≥ 100 kb, ≥ 500 kb or ≥ 1 Mb. British *A. crataegi* showed a much higher proportion of the genome in RoH (7.6% in ≥ 100 Kb, 3.2% in ≥ 500 kb and 1.6% in ≥ 1 Mb; Fig. 3C) than European *A. crataegi* (0.6% in ≥ 100 Kb, 0.2% in ≥ 500 kb, and 0.1% in ≥ 1 Mb; Fig. 3C). Additionally, the difference in all RoH categories were significantly different (≥ 100 Kb: *F_1,10_*=43.84, *p*=0.00005919; ≥ 500 kb*: F_1,10_*=28.47 *p*=0.00033; ≥ 1MB: *F_1,10_*=16.69, *p*=0.002918) with all British *A. crataegi* harbouring at least one RoH ≥ 1 Mb (Fig. S9), whereas only one of seven amongst European specimens harboured an RoH ≥ 1 Mb. In contrast, British *P. icarus* had a relatively smaller proportion (1.9%) of its genome in RoHs ≥ 100 Kb, 0.2% in ≥ 500 Kb, and none in regions ≥ 1 Mb (Fig. 3D), while French *P. icarus* had negligible RoHs in ≥ 100 Kb (0.03%) and none in ≥ 500 kb or ≥ 1 Mb. The difference in in RoH segments ≥ 100 Kb between British and French *P. icarus* was marginally significant (*t=*-3.15, *df* = 3, *p*= 0.0507; Fig. 3D).

Given a historical effective population size of 3.87 x 10^4^ for *A. crataegi*, the probability of observing at least one RoH ≥ 1 Mb in a genome of 230 Mb is 0.051. Given that all British *A. crataegi* harbour at least one RoH ≥ 1 Mb (Fig. S9) this could indicate a lower recent *N_e_* consistent with more recent inbreeding. With a historical of *N_e_* of 2.14 x 10^5^, the same probability for *P. icarus* was 0.021 respectively, consistent with this historical *N_e_*, no RoH ≥ 1 Mb were observed.

### Estimation of genetic load

In *A. crataegi* (Fig. 4A), the mean number of LoF variants was not significantly different (*F_1,10_*=2.8, *p*=0.125) between the British (17.5) and European specimens (20.2). However, the total numbers of missense and synonymous were significantly lower in British *A. crataegi* (Fig. 4B & C). In *P. icarus,* all variant classes were significantly lower in British samples (Fig. 4D-C). These are indicative of either no change (LoF) or reduction (missense and synonymous) in genetic load in British specimens.

**Figure. 4:**
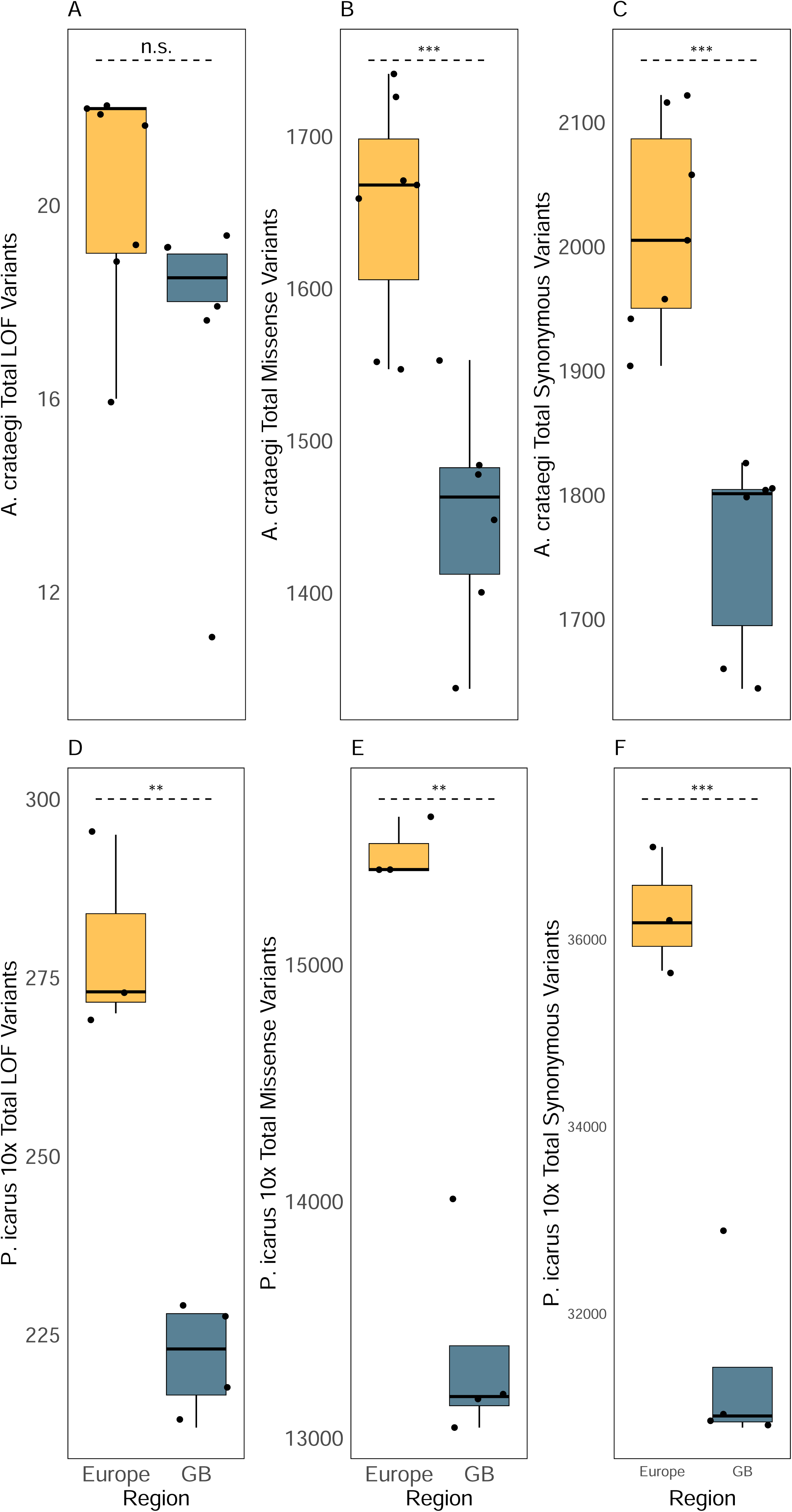
(A-C): Total LoF, Missense and Synonymous variants found using snpEff in historic A. crataegi (A.c.) specimens. (D-F): Total LoF, Missense and Synonymous variants found in P. icarus. Dashed lines show difference between Europe and GB. Significance codes: n.s. = not significant, * = p <0.05, ** = p <0.01, *** = p < 0.001. A.c: linear model with average coverage as a covariate, P.i. Welch’s two sample t-test.

Realised genetic load (deleterious variants exposed in homozygosity, assuming most deleterious variants are partially recessive) was determined by comparing the predicted variant classes in homozygous (Fig. S10A-C) state. For *A. crataegi* there was no significant increase homozygosity for LoF and missense classes, but there was a statistically significant increase in synonymous variants (*F_1,10_* = 6.37, *p*= 0.03) between the British specimens (893) and European (864) specimens. British *P. icarus* show no significant change in homozygosity across all three variant classes (Fig S11).

Genetic load in terms of derived alleles for LoF variants was not significantly different between British *A. crataegi* and mainland specimens albeit with a trend of decrease in the British specimens (Fig. 5A). However, there was a statistically significant increase in frequencies of missense and synonymous variants in British specimens relative to mainland specimens. Shared derived alleles in British *P. icarus* (Fig. 5B) exhibit a significant decrease in frequencies of missense and synonymous variants, but no statistically significant change in LoF frequency.

**Figure 5:**
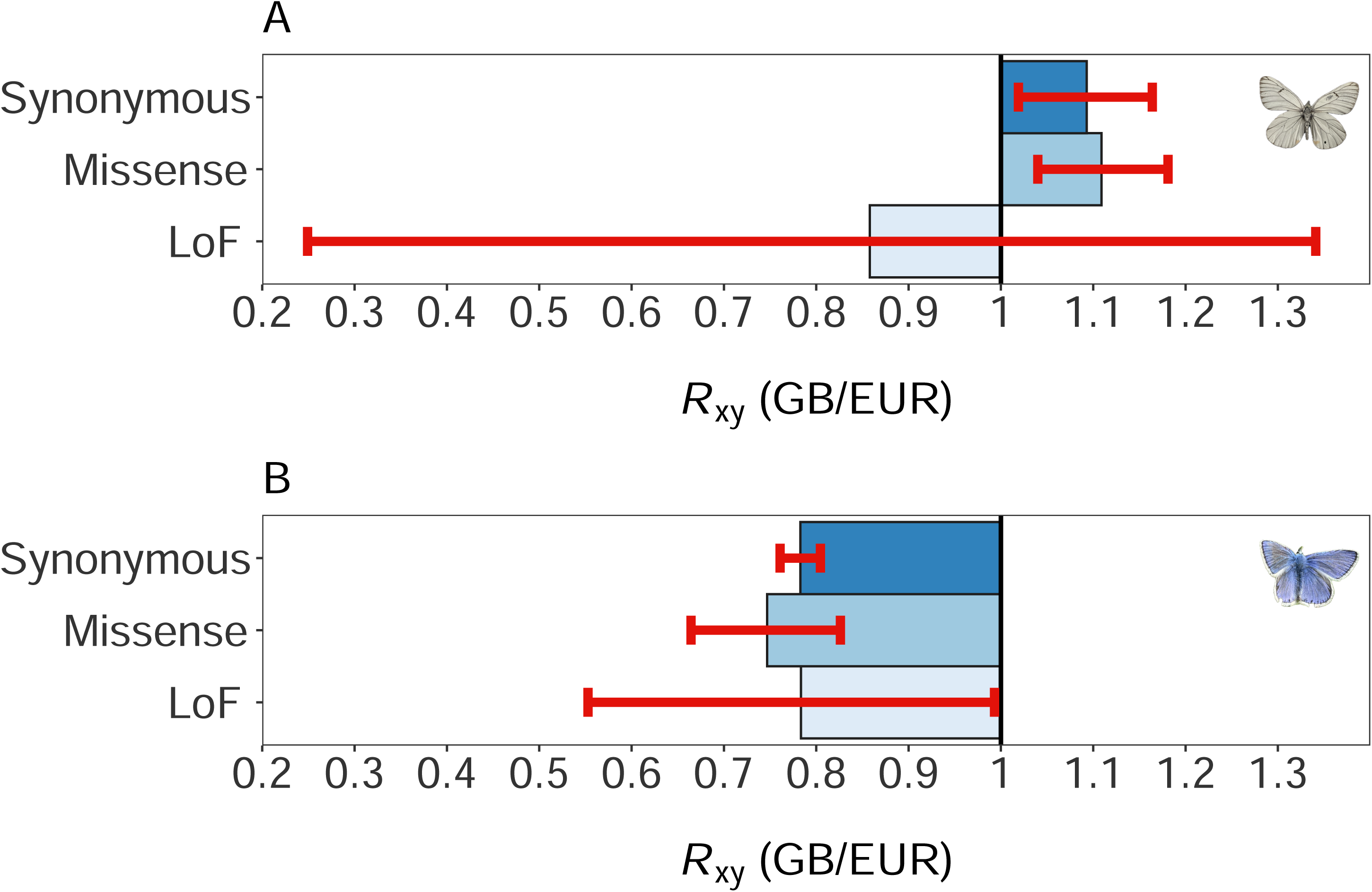
(A): Rxy ratio of shared derived alleles classified Synonymous, Missense and LoF variants in A. crataegi. Red whiskers represent 95% confidence intervals based on jackknife estimates over chromosomes. A ratio of > 1 represents an increase in frequency of alleles in population X (GB) relative to population Y (Europe), 1 reflects no change and < 1 a decrease. (B) Rxy ratio of shared derived alleles of each class in P. icarus.

## Discussion

Our extraction of historic DNA was contaminant free and we obtained short fragment lengths which decreased in length with specimen age, a degradation pattern consistent with other studies using historical DNA (e.g. Mullin *et al*., 2023). We used these historical genomes of *A. crataegi* to provide a genomic perspective on its extirpation from the British Isles in the early twentieth century and their post-glacial colonization history in Britain. We complemented our analysis with a similar approach using *P. icarus*, a butterfly with long-term stable population size within the British Isles. Our results also provide genomic insights into the extirpation of *A. crataegi* and the utility of genomic erosion metrics to identify vulnerable insect populations. This approach should be applicable to other insect species using both museum and contemporary specimens. Comparative data on genomic erosion in threatened, stable and expanding species, but also those with different life histories could improve the predictive capacity for using metrics of genome erosion for identifying and managing threatened populations and species (Bortoluzzi *et al*., 2023; Kyriazis *et al*., 2023; Webster *et al*., 2023).

### Different demographic histories of A. crataegi and P. icarus in Britain

Our population structure analysis (Fig. 1B-C) and PSMC-based demographic reconstruction (Fig. 2) suggest that British populations of both *A. crataegi* and *P. icarus* are genetically distinct from their continental European counterparts. However, there was evidence of strong structuring within the British populations (Fig. 1B-C). Structuring apparent within the British specimens could result from recent gene flow with European mainland populations or from sequential founder events and isolation by distance following colonisation and subsequent founder events or a combination of these two hypotheses. Gene flow and/or sequential founder events are a likely explanation for the structuring observed in *P. icarus* as has been suggested by Arif *et al*., (2021). However, for *A. crataegi*, the consistency of demographic reconstructions (Fig. 2 and Fig S4), genome-wide heterozygosity (Fig. 2A), RoHs (Fig. 2C) and genetic load (Fig. 3A-C) amongst the British specimens, do not favour a hypothesis of gene flow or admixture from the continent.

The post-glacial colonisation of the British Isles by *A. crataegi* and *P. icarus* are likely to have been different (Fig. 2), based on their differing resource requirements (Middleton-Welling *et al*., 2020) and climate tolerances (Dennis 1992). *A. crataegi* uses wood edge shrub and tree species as hostplants and is primarily a species associated with grassland edge/scrub matrices and requires relatively warm summer temperatures whilst *P. icarus* is associated with pioneer or persistent short grassland locations in which its hostplants occur (Howe *et al*., 2007; De Keyser, *et al*., 2012), and is tolerant of lower temperatures than *A. crataegi* (Dennis, 1992). Conditions in Britain and the adjacent European mainland during the Younger Dryas would not have permitted the occurrence of *A. crataegi* and it is most likely to have colonized Britain in the short period from the late Boreal to the elimination of channel land c.11.5 – 8.5 Kya (Dennis, 1992), probably arriving from mid-European latitudes. By contrast, *P. icarus* may have persisted in sheltered southern valleys further north than *A. crataegi* during the Younger Dryas, possibly in what is currently southern Britain, northern France and possibly channel land, having originally reached southern Britain as early as the late Glacial period (c.14.5 – 13.5 Kya) (Dennis 1992). Thus, gene flow and admixture during the colonization period is more likely for *P. icarus* than for *A. crataegi*. For *P. icarus* it is also possible that there were at least two colonization periods, with good evidence for mixing of different lineages from distinct glacial refugia (Arif *et al.,* 2021). Furthermore, admixture and gene flow are likely to be higher in *P. icarus* because of its multivoltine life-history, compared to the univoltine strategy of *A. crataegi* which reduces the number of generations over which these can occur over any time period.

Our PSMC-based demographic reconstructions and species life histories explain changes of effective population sizes and the spatial structuring of populations. Both species underwent decreases of *N_e_* (Fig. 2 and Fig. S4) around the passing of the Eemian interglacial period (∼125 Kya) until around the last glacial maximum (LGM, ∼20 Kya). After this time (∼15 Kya) British *A. crataegi* exhibit a marked reduction in *N_e_*, consistent with a genetic bottleneck associated with a founder event and earlier northward spread from southern glacial refugia. For British *A. crataegi* this *N_e_* (≈ 10^4^) is a magnitude smaller than the commonly observed sizes of 10^5^ or greater reported in butterflies (Mackintosh *et al*., 2019; Ebdon *et al*., 2021), but is consistent with PSMC-based estimates of several other British Lepidoptera (Bortoluzzi *et al*., 2023). At the same time, European *A. crataegi* show a drastic increase in *N_e_* presumably because of mixing of distinct populations from previously isolated glacial refugia. (Hewitt, 1999; Hinojosa *et al.,* 2019). It is likely that colonization occurred over a relatively short time-period, and possibly by individuals from few source populations. Our results also indicate a reduction of *N_e_* and a bottleneck for *P. icarus* with divergence from mainland European populations just before the LGM but also note that inference of the precise timing of the *P. icarus* bottleneck is dependent on the number of generations per year which may vary from 1 to 3 (Fig. S6). A bottleneck around the time of colonization of Britain at the end of the last glacial would be consistent with the interpretation of allozyme data by De Keyser *et al*. (2012), however it is also possible that the lineages of *P. icarus* that colonised Britain never recovered from pre-LGM reductions in *N_e_* , which may be the case for rapidly colonising species. The discrepancy in the timings of our demographic reconstructions (Fig. 2) and the exact timing of the colonisation of the British Isles by these species may also result from bias in the PSMC method. Simulations by Bortoluzzi *et al*. (2023) suggest, that PSMC reconstructions for species with small genomes and a high ratio of recombination to mutation rate, may reflect linked selection rather than true demographic history. We cannot rule this out for either species, and this bias could potentially explain differences in the demographic timing (along with uncertainty in generation time for *P. icarus*). We note that our PSMC-based trajectories recapitulate changes in *N_e_* that are largely consistent with major climatic events of the last 10-125 Kya. Finally, we note that absolute *N_e_* of British *A. crataegi* (≈ 10^4^ ) and *P. icarus* (≈ 10^5^ ) are a function of their species-specific *N_e_,* which could be considerably smaller in *A. crataegi*, given that genetic diversity correlates negatively with body size and positively with genome size in Lepidoptera (Mackintosh *et al*., 2019).

### Genomic symptoms of the demise of A. crataegi in Britain

The last recorded sighting of *A. crataegi* in Britain dates from *c.* 1925 (Pratt, 1983). However, by this time, the species was thought to be limited to the south-eastern regions of Kent and had already disappeared from other parts of its range as early as 1880 (Pratt, 1983 and references therein).

Consistent with these observations, three of 13 of our British samples, samples collected in 1888, 1908 and 1924 , appear to be of European origin in terms of population structure, demographic history, and genomic erosion. These specimens may have been captive-bred or wild-caught European (potentially from imported German larvae (Pratt, 1983)) stock released into British habitats or potential migrants.

There was a reduction in genome-wide heterozygosity in British specimens of both species, but reductions (compared to European populations) in both the extirpated *A. crataegi* and the extant and stable *P. icarus* were roughly comparable (17.9 and 14.5%, respectively). As a comparison, the observed reduction in genome-wide heterozygosity in the extinct *Glaucopsyche xerces* (Xerces Blue butterfly; de-Dios *et al*., 2023) was 22%, hence greater than what we observed for British *A. crataegi*. However, the former was calculated in comparison to a sister species (*Glaucopsyche lygdamus*) and hence it is unclear whether these estimates of reductions in genome-wide heterozygosity are strictly comparable.

We saw significant increases in homozygous regions of small (≥ 100 Kb), intermediate (≥ 500 Kb) and large (≥ 1 Mb) variety (Fig. 3C, Fig. S9) in British *A. crataegi*.. In contrast, no category of F_RoH_ was significantly different between British and continental European *P. icarus* suggesting this population likely does not suffer from inbreeding. The probability of observing large (≥ 1 Mb) RoHs, given the PSMC-estimated historical *N_e_* for both *A. crategi* and *P. icarus* were of the same magnitude (0.051 and 0.021, respectively), yet such RoH were more prevalent in *A. crategi*. The presence of the large RoHs raise the possibility of more recent inbreeding, consistent with a spasmodic rather than instant decline of the species in the 19^th^ century. It is important to note that the proportions of genome in RoH detected in *A. crataegi* genomes are likely an underestimate due to limited coverage in the museum specimens (Table S1), as the fraction of detectable RoHs is a function of genome coverage (Meyermans *et al*., 2020).

Accumulation of mildly or weakly deleterious variants, solely due to drift after a bottleneck, is expected to lead to decrease in the total number of such variants but increase in homozygosity across all variant classes (Dussex *et al.,* 2023). However, purging of deleterious variants through purifying selection should lead to a decrease in strongly deleterious variants (LoF) in both homozygous and heterozygous state, whereas homozygosity of mildly or weakly deleterious variants can still increase and inflate the genetic load. The accumulation of genetic load in small populations has been observed in many vertebrate populations (see references in Dussex *et al*. 2023) and in the Xerces blue butterfly (de-Dios *et al*., 2023). However, there has also been evidence documentation of purging of the most deleterious variants and accumulation of less deleterious variants in the same system in Alpine Ibex (Grossen *et al*., 2020) and Indian tigers (Khan *et al*., 2021) and is supported by simulations (Dussex *et al*., 2023). Although we observed no clear evidence of purging of LoF in British *A. crataegi* (Fig. 4A, Fig. S10A and D, Fig. 5A), we did see an increase in shared derived allele frequency in the missense and synonymous categories which may have contributed to extirpation by reducing fitness and population viability. Additionally, we also observed a significant increase in realised load but this was limited to only synonymous variants. Although synonymous variants are normally classified as neutral, they can be weakly deleterious under codon bias and Lepidoptera do exhibit an A/T codon usage bias (Näsvall *et al*., 2023). In contrast, *P. icarus* displayed a decrease in realised load (Fig. S11) and a decrease in frequencies of shared derived alleles of the missense and synonymous categories (Fig. 5B), suggesting these populations may have been less influenced by genetic drift. These results raise possibility that the accumulation of mildly and weakly deleterious mutations contributed to the decline of *A. crataegi* in Britain by triggering a mutational meltdown (Lynch *et al*., 1995b). However, it should be noted that the actual fitness effects including selection and dominance coefficients of the annotated variants remains unknown. Moreover, given our limited sample sizes and lack of statistical significance in some cases, our results on genetic load should be interpreted with caution. Finally, given that the annotation for *P. icarus* (Lohse *et al*., 2023) is based on a single male from Scotland, our predictions of phenotypic impact of variants for this species may suffer from annotation bias, for example, by reducing the number of predicted LoF and missense variants.

Overall, we provide compelling evidence that *A. crataegi* suffered a severe bottleneck during the colonisation of the British Isles. Based on our analysis it is likely that *N_e_* likely remained low following the bottleneck and further reduced by more recent inbreeding, leading to the accumulation of weak or mild deleterious mutations and potentially an increase in realised load, which may have further exacerbated population persistence in the short term. Interestingly, two other butterfly species, *Lycaena dispar* (Large copper butterfly) and *Cyaniris semiargus* (Mazarine blue butterfly) were also extirpated in Britain between the mid to late 19^th^ century. Like *A. crataegi*, both species expressed single broods in Great Britain, but unlike *A. crataegi* both species are smaller with potentially larger genomes and hence are expected to have higher ancestral *N_e_* (Mackintosh *et al*., 2019), however as yet nothing is known about their demographic history in Britain. Hence, it would be instructive to examine whether genomes of these extirpated populations exhibit similar symptoms of genomic erosion observed in *A. crataegi* and *G. xerces*.

### Metrics of genomic erosion for identifying vulnerable insect populations

Conservation genomics could be an extremely helpful tool in identifying populations or species at risk of extinction or extirpation. This could be particularly relevant for invertebrates, whereas opposed to vertebrates, reductions in census population size may not be easily monitored (Cardoso *et al*., 2011; Cardoso and Leather, 2019; Didham *et al.,* 2020). Our results suggest that a consistently small *N_e_* on the order of ≈10^4^, inbreeding in the recent past and accumulation of deleterious variants could be viable indicators of population decline in species with generally large *N_e_*. Indeed, similar signals of genomic erosion have also been detected in the genomes of the extinct *G. xerces* (de-Dios *et al*., 2023).

To evaluate how genomic erosion translates into population persistence or extinction we require further empirical studies of not only declining but stable and expanding populations as well. Isolated populations on islands, including the British Isles, offer an excellent system to study the genomics of populations with contrasting or unknown fates (e.g. Bortoluzzi *et al*., 2023; Kyriazis *et al*., 2023).

Museum collections can provide a substantial and unequivocal source of threatened populations (e.g., samples from extirpated or extinct populations). Recent advances in resequencing of museum samples (Korlević *et al*., 2021, Mullin *et al*., 2023), bioinformatic tools (e.g., Korneliussen *et al*., 2014, Kutschera *et al.,* 2022) and the availability of high-quality genomes with annotations (Rhie *et al*., 2021; Formenti *et al.,* 2022; DToL, 2022) are providing much needed resources for such work.

Although challenges remain, for example, evaluating and interpreting fitness effects of deleterious mutations (de Valles-Ibáñez *et al*., 2016; Robinson *et al*., 2023), we expect museomics in combination with contemporary samples will expand our understanding of the genomics of extinction but also improve feasibility to monitor and manage threatened insect populations (Díez-del-Molino *et al.,* 2018; Jensen *et al*., 2022; Bortoluzzi *et al*., 2023).

## Acknowledgements

This work was supported by a Nigel Groome PhD Studentship (Department of Biological and Medical Sciences, Oxford Brookes University) to RW and a pilot funding grant from the Centre for Functional Genomics, Oxford Brookes University to SA. We would like to thank the Oxford University Natural History Museum (OUMNH) and the Natural History Museum, London (NHMUK) for permission to obtain the Black-veined white museum samples. We would also like to thank Joe Middleton-Welling for obtaining the samples for a pilot study from the OUMNH and the Darwin Tree of Life for the Black-veined White genome assembly. We would also like to thank Konrad Lohse for help in calculating coalescent probabilities. We thank Laura Lukens for helpful comments on improving the manuscript.

## Author Contributions

TGS, SA and RW designed the study. JH and GM helped procure museum specimens. TGS, SA and RW collected data. RW completed all molecular work with supervision and support from KH, CB and SA. RW analysed all the data with the help of SA. SA and RW wrote the paper with input from all authors.

## Data Availability Statement

Raw sequence data is available under project accessing PRJEB70473 (for *A. crataegi*) and PRJEB70726 (for *P. icarus*) on the European Nucleotide Archive (ENA). Configuration files for GenErode and additional script files are available at https://github.com/rmwhitla/BVWpaper. Protocol for library preparation of historical samples is available at https://www.protocols.io/view/library-prep-for-cut-run-with-nebnext-ultra-ii-dna-bwgjpbun

## Notes

### Competing Interest Statement

The authors have declared no competing interest.

### Summary of Updates

The following edits and corrections were made for this version: 1.)The PSMC results presented in previous versions were incorrect and the methods were incomplete. We had also varied the maximum coalescent time (-t) parameter when running PSMC to avoid overfitting (as suggested by the author: https://github.com/lh3/psmc) but this had not been stated in the Methods previously. This is now stated in the Methods section and the correct PSMC results in Figure 2 and estimates of historical Ne have been incorporated in the ms. This change results in very minor changes to the Figure 2 and historical Ne and have no impact on our conclusions. 2.)The FRoH values were erroneously reported with a systematic bias that inflated all values. The correct FRoH values are now presented in the Results Section and have also been updated in Figures 3C and D. Since the error was a systematic bias, it did not influence the outcome of the statistical analysis and has no impact on our conclusions. However, the correct FRoH values are similar in magnitude to those observed in Glaucopsyche xerces (https://doi.org/10.7554/eLife.87928.2) and we have removed a sentence in the discussion that stated our FRoH were greater in magnitude than those observed for Glaucopsyche xerces 3.)Some points were omitted due limits of the axis in Figure 3A. The axis limits have been expanded so that all points are displayed in Figure 3A 4.)There were some duplicate points on the extremities of the boxplot distributions in Figure 4. These duplicate points have now been removed. 5.)Some minor edits have been made to the text, in response to this open review, to help improve our writing and clarity. We acknowledge the author for their review in the Acknowledgements.

## References

Aken, B.L., Achuthan, P., Akanni, W., Amode, M.R., Bernsdorff, F., Bhai, J., Billis, K., Carvalho-Silva, D., Cummins, C., Clapham, P., Gil, L., Girón, C.G., Gordon, L., Hourlier, T., Hunt, S.E., Janacek, S.H., Juettemann, T., Keenan, S., Laird, M.R., Lavidas, I., Maurel, T., McLaren, W., Moore, B., Murphy, D.N., Nag, R., Newman, V., Nuhn, M., Ong, C.K., Parker, A., Patricio, M., Riat, H.S., Sheppard, D., Sparrow, H., Taylor, K., Thormann, A., Vullo, A., Walts, B., Wilder, S.P., Zadissa, A., Kostadima, M., Martin, F.J., Muffato, M., Perry, E., Ruffier, M., Staines, D.M., Trevanion, S.J., Cunningham, F., Yates, A., Zerbino, D.R., Flicek, P. (2017). Ensembl 2017. Nucleic Acids Research, 45, D635–D642. 10.1093/nar/gkw1104

Andrews, S., (2010). FastQC: A Quality Control Tool for High Throughput Sequence Data. Available online at: http://www.bioinformatics.babraham.ac.uk/projects/fastqc/.

Arif, S., Gerth, M., Hone-Millard, W.G., Nunes, M.D.S., Dapporto, L., Shreeve, T.G. (2021). Evidence for multiple colonisations and *Wolbachia* infections shaping the genetic structure of the widespread butterfly *Polyommatus icarus* in the British Isles. Molecular Ecology, 30, 5196–5213. 10.1111/mec.16126

Barnosky, A.D., Matzke, N., Tomiya, S., Wogan, G.O.U., Swartz, B., Quental, T.B., Marshall, C., McGuire, J.L., Lindsey, E.L., Maguire, K.C., Mersey, B., Ferrer, E.A. (2011). Has the Earth’s sixth mass extinction already arrived? Nature, 471, 51–57. 10.1038/nature09678

Biscarini, F., Cozzi, P., Gaspa, G., Marras, G. (2018). detectRUNS: Detect runs of homozygosity and runs of heterozygosity in diploid genomes [WWW Document]. URL https://cran.r-project.org/web/packages/detectRUNS/index.html

Bortoluzzi, C., Wright, C. J., Lee, S., Cousins, T., Genez, T. A. L., Thybert, D., Martin, F. J., Haggerty, L., The Darwin Tree of Life Project Consortium, Blaxter, M., Durbin, R. (2023). Lepidoptera genomics based on 88 chromosomal reference sequences informs population genetic parameters for conservation. bioRxiv 2023.04.14.536868. [PREPRINT] 10.1101/2023.04.14.536868

Bosse, M., van Loon, S. (2022). Challenges in quantifying genome erosion for conservation. Frontiers in Genetics, 13, 960958. 10.3389/fgene.2022.960958

Burns, F., Mordue, S., al Fulaij, N., Boersch-Supan, P. H., Boswell, J., Boyd, R. J., Bradfer-Lawrence, T., de Ornellas, P., de Palma, A., de Zylva, P., Dennis, E. B., Foster, S., Gilbert, G., Halliwell, L., Hawkins, K., Kaysom, K. A., Holland, M. M., Hughes, J., Jackson, A. C., Mancini, F., Mathews, F., McQuatters-Gollop, A., Noble, D. G., O’Brien, D., Pescott, O. L., Purvis, A., Simkin, J., Smith, A., Stanbury, A. J., Villemot, J., Walker, K. J., Walton, P., Webb, T. J., Williams, K., Wilson, R., Gregory, R. D. (2023). State of Nature 2023, the State of Nature Partnership. Available at: www.stateofnature.org.uk

Cardoso, P., Erwin, T.L., Borges, P.A.V., New, T.R. (2011). The seven impediments in invertebrate conservation and how to overcome them. Biological Conservation, 144, 2647–2655. 10.1016/j.biocon.2011.07.024

Cardoso, P., Leather, S.R. (2019). Predicting a global insect apocalypse. Insect Conservation and Diversity, 12, 263–267. 10.1111/icad.12367

Ceballos, G., Ehrlich, P.R., Barnosky, A.D., García, A., Pringle, R.M., Palmer, T.M. (2015). Accelerated modern human-induced species losses: Entering the sixth mass extinction. Science Advances, 1, e1400253. 10.1126/sciadv.1400253

Ceballos, F.C., Joshi, P.K., Clark, D.W., Ramsay, M., Wilson, J.F. (2018). Runs of homozygosity: windows into population history and trait architecture. Nature Reviews Genetics, 19, 220–234. 10.1038/nrg.2017.109

Charlesworth, D., Willis, J.H. (2009). The genetics of inbreeding depression. Nature Reviews Genetics, 10, 783–796. 10.1038/nrg2664

Chen, S., Zhou, Y., Chen, Y., Gu, J. (2018). fastp: an ultra-fast all-in-one FASTQ preprocessor. Bioinformatics, 34, i884–i890. 10.1093/bioinformatics/bty560

Cingolani, P., Platts, A., Wang, L.L., Coon, M., Nguyen, T., Wang, L., Land, S.J., Lu, X., Ruden, D.M. (2012). A program for annotating and predicting the effects of single nucleotide polymorphisms, SnpEff. Fly (Austin), 6, 80–92. 10.4161/fly.19695

Cowie, R.H., Bouchet, P., Fontaine, B. (2022). The Sixth Mass Extinction: fact, fiction or speculation? Biological Reviews, 97, 640–663. 10.1111/brv.12816

Dale, C.W. (1887). Query regarding *Aporia crataegi*. The Entomologist’s Monthly Magazine, 23, 214.

Danecek, P., Bonfield, J.K., Liddle, J., Marshall, J., Ohan, V., Pollard, M.O., Whitwham, A., Keane, T., McCarthy, S.A., Davies, R.M., Li, H. (2021). Twelve years of SAMtools and BCFtools. Gigascience, 10, giab008. 10.1093/gigascience/giab008

Darwin Tree of Life Project Consortium. (2022). Sequence locally, think globally: The Darwin Tree of Life Project. Proceedings of the National Academy of Sciences of the United States of America, 119, e2115642118. 10.1073/pnas.2115642118

de-Dios, T., Fontsere, C., Renom, P., Stiller, J., Llovera, L., Uliano-Silva, M., Sánchez-Gracia, A., Wright, C., Lizano, E., Caballero, B., Navarro, A., Civit, S., Robbins, R.K., Blaxter, M., Marquès-Bonet, T., Vila, R., Lalueza-Fox, C. (2023). Whole-genomes from the extinct Xerces Blue butterfly can help identify declining insect species. eLife, 12. 10.7554/eLife.87928

De Keyser, R., Shreeve, T.G., Breuker, C.J., Hails, R.S., Schmitt, T. (2012). *Polyommatus icarus* butterflies in the British Isles: evidence for a bottleneck. Biological Journal of the Linnean Society, 107, 123–136. 10.1111/j.1095-8312.2012.01925.x

de Valles-Ibáñez, G., Hernandez-Rodriguez, J., Prado-Martinez, J., Luisi, P., Marquès-Bonet, T., Casals, F. (2016). Genetic Load of Loss-of-Function Polymorphic Variants in Great Apes. Genome Biology and Evolution, 8, 871–877. 10.1093/gbe/evw040

Dennis, R.L.H. (1992). An evolutionary history of British butterflies. In Dennis, R.L.H. (Ed.) The Ecology of Butterflies in Britain. Oxford, Oxford University Press, pp. 217–245.

Dennis, R.L.H., Shreeve, T.G., Van Dyck, H. (2003). Towards a Functional Resource-Based Concept for Habitat: A Butterfly Biology Viewpoint. Oikos, 102, 417–426. 10.1034/j.1600-0579.2003.12492.x

DeWoody, J.A., Harder, A.M., Mathur, S., Willoughby, J.R. (2021). The long-standing significance of genetic diversity in conservation. Molecular Ecology, 30, 4147–4154. 10.1111/mec.16051

Didham, R.K., Basset, Y., Collins, C.M., Leather, S.R., Littlewood, N.A., Menz, M.H.M., Müller, J., Packer, L., Saunders, M.E., Schönrogge, K., Stewart, A.J.A., Yanoviak, S.P., Hassall, C. (2020). Interpreting insect declines: seven challenges and a way forward. Insect Conservation and Diversity, 13, 103–114. 10.1111/icad.12408

Díez-del-Molino, D., Sánchez-Barreiro, F., Barnes, I., Gilbert, M.T.P., Dalén, L. (2018). Quantifying Temporal Genomic Erosion in Endangered Species. Trends in Ecology & Evolution, 33, 176–185. 10.1016/j.tree.2017.12.002

Dincă, V., Montagud, S., Talavera, G., Hernández-Roldán, J., Munguira, M.L., García-Barros, E., Hebert, P.D.N., Vila, R. (2015). DNA barcode reference library for Iberian butterflies enables a continental-scale preview of potential cryptic diversity. Scientific Reports, 5, 12395. 10.1038/srep12395

Dussex, N., van der Valk, T., Morales, H.E., Wheat, C.W., Díez-del-Molino, D., von Seth, J., Foster, Y., Kutschera, V.E., Guschanski, K., Rhie, A., Phillippy, A.M., Korlach, J., Howe, K., Chow, W., Pelan, S., Mendes Damas, J.D., Lewin, H.A., Hastie, A.R., Formenti, G., Fedrigo, O., Guhlin, J., Harrop, T.W.R., Le Lec, M.F., Dearden, P.K., Haggerty, L., Martin, F.J., Kodali, V., Thibaud-Nissen, F., Iorns, D., Knapp, M., Gemmell, N.J., Robertson, F., Moorhouse, R., Digby, A., Eason, D., Vercoe, D., Howard, J., Jarvis, E.D., Robertson, B.C., Dalén, L. (2021). Population genomics of the critically endangered kākāpō. Cell Genomics, 1, 100002. 10.1016/j.xgen.2021.100002

Dussex, N., Morales, H.E., Grossen, C., Dalén, L., Oosterhout, C. van. (2023). Purging and accumulation of genetic load in conservation. Trends in Ecology & Evolution, 38, 961–969. 10.1016/j.tree.2023.05.008

Ebdon, S., Laetsch, D. R., Dapporto, L., Hayward, A., Ritchie, M. G., Dincӑ, V., Vila, R., & Lohse, K. (2021). The Pleistocene species pump past its prime: Evidence from European butterfly sister species. Molecular Ecology, 30, 3575–3589. 10.1111/mec.15981

Ebdon, S., Mackintosh, A., Lohse, K., Hayward, A., Arif, S., Whitla, R. (2022). The genome sequence of the black-veined white butterfly, Aporia crataegi (Linnaeus, 1758) [version 1; peer review: 1 approved, 1 not approved]. Wellcome Open Research 7:81. 10.12688/wellcomeopenres.17709.1

Formenti, G., Theissinger, K., Fernandes, C., Bista, I., Bombarely, A., Bleidorn, C., Ciofi, C., Crottini, A., Godoy, J.A., Höglund, J., Malukiewicz, J., Mouton, A., Oomen, R.A., Paez, S., Palsbøll, P.J., Pampoulie, C., Ruiz-López, María J., Svardal, H., Theofanopoulou, C., de Vries, J., Waldvogel, A.-M., Zhang, Guojie, Mazzoni, C.J., Jarvis, E.D., Bálint, M., Formenti, G., Theissinger, K., Fernandes, C., Bista, I., Bombarely, A., Bleidorn, C., Čiampor, F., Ciofi, C., Crottini, A., Godoy, J.A., Hoglund, J., Malukiewicz, J., Mouton, A., Oomen, R.A., Paez, S., Palsbøll, P., Pampoulie, C., Ruiz-López, María José, Svardal, H., Theofanopoulou, C., de Vries, J., Waldvogel, A.-M., Zhang, Goujie, Mazzoni, C.J., Jarvis, E., Bálint, M., Aghayan, S.A., Alioto, T.S., Almudi, I., Alvarez, N., Alves, P.C., Amorim, I.R., Antunes, A., Arribas, P., Baldrian, P., Berg, P.R., Bertorelle, G., Böhne, A., Bonisoli-Alquati, A., Boštjančić, L.L., Boussau, B., Breton, C.M., Buzan, E., Campos, P.F., Carreras, C., Castro, L.Fi., Chueca, L.J., Conti, E., Cook-Deegan, R., Croll, D., Cunha, M.V., Delsuc, F., Dennis, A.B., Dimitrov, D., Faria, R., Favre, A., Fedrigo, O.D., Fernández, R., Ficetola, G.F., Flot, J.-F., Gabaldón, T., Galea Agius, D.R., Gallo, G.R., Giani, A.M., Gilbert, M.T.P., Grebenc, T., Guschanski, K., Guyot, R., Hausdorf, B., Hawlitschek, O., Heintzman, P.D., Heinze, B., Hiller, M., Husemann, M., Iannucci, A., Irisarri, I., Jakobsen, K.S., Jentoft, S., Klinga, P., Kloch, A., Kratochwil, C.F., Kusche, H., Layton, K.K.S., Leonard, J.A., Lerat, E., Liti, G., Manousaki, T., Marques-Bonet, T., Matos-Maraví, P., Matschiner, M., Maumus, F., Mc Cartney, A.M., Meiri, S., Melo-Ferreira, J., Mengual, X., Monaghan, M.T., Montagna, M., Mysłajek, R.W., Neiber, M.T., Nicolas, V., Novo, M., Ozretić, P., Palero, F., Pârvulescu, L., Pascual, M., Paulo, O.S., Pavlek, M., Pegueroles, C., Pellissier, L., Pesole, G., Primmer, C.R., Riesgo, A., Rüber, L., Rubolini, D., Salvi, D., Seehausen, O., Seidel, M., Secomandi, S., Studer, B., Theodoridis, S., Thines, M., Urban, L., Vasemägi, A., Vella, A., Vella, N., Vernes, S.C., Vernesi, C., Vieites, D.R., Waterhouse, R.M., Wheat, C.W., Wörheide, G., Wurm, Y., Zammit, G. (2022). The era of reference genomes in conservation genomics. Trends in Ecology & Evolution, 37, 197–202. 10.1016/j.tree.2021.11.008

Fox, R., Dennis, E.B., Brown, A.F., Curson, J. (2022). A revised Red List of British butterflies. Insect Conservation and Diversity, 15, 485–495. 10.1111/icad.12582

Gomulkiewicz, R., Holt, R.D. (1995). When does Evolution by Natural Selection Prevent Extinction? Evolution, 49, 201–207. 10.2307/2410305

Grossen, C., Guillaume, F., Keller, L.F., Croll, D. (2020). Purging of highly deleterious mutations through severe bottlenecks in Alpine ibex. Nature Communications, 11, 1001. 10.1038/s41467-020-14803-1

Hewitt, G.M. (1999). Post-glacial re-colonization of European biota. Biological Journal of the Linnean Society, 68, 87–112. 10.1111/j.1095-8312.1999.tb01160.x

Hill, G.M., Kawahara, A.Y., Daniels, J.C., Bateman, C.C., Scheffers, B.R. (2021). Climate change effects on animal ecology: butterflies and moths as a case study. Biological Reviews of the Cambridge Philosophical Society, 96, 2113–2126. 10.1111/brv.12746

Hinojosa, J.C., Koubínová, D., Szenteczki, M.A., Pitteloud, C., Dincă, V., Alvarez, N., Vila, R. (2019). A mirage of cryptic species: Genomics uncover striking mitonuclear discordance in the butterfly *Thymelicus sylvestris*. Molecular Ecology, 28, 3857–3868. 10.1111/mec.15153

Howe, P.D., Bryant, S.R., Shreeve, T.G. (2007). Predicting body temperature and activity of adult *Polyommatus icarus* using neural network models under current and projected climate scenarios. Oecologia, 153(4), 857–69. 10.1007/s00442-007-0782-3

Hudson, R. R. (1990). Gene Genealogies and the coalescent process. In D. Futuyma and J. Antonovics (Eds.), Oxford Surveys in Evolutionary Biology, Vol. 7. Oxford University Press, New York, pp. 1–44.

IPBES, 2019. Summary for policymakers of the global assessment report on biodiversity and ecosystem services. IPBES Plenary at its seventh session (IPBES 7, Paris, 2019) 29 April - 4 May 2019. 10.5281/zenodo.3553579

Jensen, E.L., Díez-del-Molino, D., Gilbert, M.T.P., Bertola, L.D., Borges, F., Cubric-Curik, V., de Navascués, M., Frandsen, P., Heuertz, M., Hvilsom, C., Jiménez-Mena, B., Miettinen, A., Moest, M., Pečnerová, P., Barnes, I., Vernesi, C. (2022). Ancient and historical DNA in conservation policy. Trends in Ecology & Evolution, 37, 420–429. 10.1016/j.tree.2021.12.010

Jiggins, C.D., Mavarez, J., Beltrán, M., McMillan, W.O., Johnston, J.S., Bermingham, E. (2005) A Genetic Linkage Map of the Mimetic Butterfly *Heliconius melpomene*, Genetics, 171(2), 557– 570, 10.1534/genetics.104.034686

Jónsson, H., Ginolhac, A., Schubert, M., Johnson, P.L.F., Orlando, L. (2013). mapDamage2.0: fast approximate Bayesian estimates of ancient DNA damage parameters. Bioinformatics, 29, 1682–1684. 10.1093/bioinformatics/btt193

Karam-Gemael, M., Decker, P., Stoev, P., Marques, M.I., Jr, A.C. (2020). Conservation of terrestrial invertebrates: a review of IUCN and regional Red Lists for Myriapoda. ZooKeys, 930, 221–229. 10.3897/zookeys.930.48943

Kardos, M., Taylor, H.R., Ellegren, H., Luikart, G., Allendorf, F.W. (2016). Genomics advances the study of inbreeding depression in the wild. Evolutionary Applications, 9, 1205–1218. 10.1111/eva.12414

Keightley, P.D., Pinharanda, A., Ness, R.W., Simpson, F., Dasmahapatra, K.K., Mallet, J., Davey, J.W., Jiggins, C.D. (2015). Estimation of the Spontaneous Mutation Rate in *Heliconius melpomene*. Molecular Biology and Evolution, 32, 239–243. 10.1093/molbev/msu302

Khan, A., Patel, K., Shukla, H., Viswanathan, A., van der Valk, T., Borthakur, U., Nigam, P., Zachariah, A., Jhala, Y.V., Kardos, M., Ramakrishnan, U. (2021). Genomic evidence for inbreeding depression and purging of deleterious genetic variation in Indian tigers. Proceedings of the National Academy of Sciences, 118, e2023018118. 10.1073/pnas.2023018118

Kim, T., Han, Y.-G., Kwon, O., Cho, Y. (2015). Changes in *Aporia crataegi*’s potential habitats in accordance with climate changes in the northeast Asia. Journal of Ecology and Environment, 38, 15–23. 10.5141/ecoenv.2015.002

Korlević, P., McAlister, E., Mayho, M., Makunin, A., Flicek, P., Lawniczak, M.K.N. (2021). A Minimally Morphologically Destructive Approach for DNA Retrieval and Whole-Genome Shotgun Sequencing of Pinned Historic Dipteran Vector Species. Genome Biology and Evolution, 13, evab226. 10.1093/gbe/evab226

Korneliussen, T.S., Albrechtsen, A., Nielsen, R. (2014). ANGSD: Analysis of Next Generation Sequencing Data. BMC Bioinformatics, 15, 356. 10.1186/s12859-014-0356-4

Kutschera, V.E., Kierczak, M., van der Valk, T., von Seth, J., Dussex, N., Lord, E., Dehasque, M., Stanton, D.W.G., Khoonsari, P.E., Nystedt, B., Dalén, L., Díez-del-Molino, D. (2022). GenErode: a bioinformatics pipeline to investigate genome erosion in endangered and extinct species. BMC Bioinformatics, 23, 228. 10.1186/s12859-022-04757-0

Kyriazis, C.C., Beichman, A.C., Brzeski, K.E., Hoy, S.R., Peterson, R.O., Vucetich, J.A., Vucetich, L.M., Lohmueller, K.E., Wayne, R.K. (2023). Genomic Underpinnings of Population Persistence in Isle Royale Moose. Molecular Biology and Evolution, 40, msad021. 10.1093/molbev/msad021

Lenth, R. (2023). emmeans: Estimated Marginal Means, aka Least-Squares Means. R package version 1.8.7. https://github.com/rvlenth/emmeans

Li, H., Durbin, R. (2009). Fast and accurate short read alignment with Burrows–Wheeler transform. Bioinformatics, 25, 1754–1760. 10.1093/bioinformatics/btp324

Li, H., Durbin, R. (2011). Inference of human population history from individual whole-genome sequences. Nature, 475, 493–496. 10.1038/nature10231

Liu, N. (2019). Library Prep for CUT&RUN with NEBNext® Ultra™ II DNA Library Prep Kit for Illumina® (E7645). protocols.io 10.17504/protocols.io.wvgfe3w

Lohse, K. (2023). The genome sequence of the Common Blue, Polyommatus icarus (Rottemburg, 1775). [version 1; peer review: 1 approved]. *Wellcome Open Research* 8:72. 10.12688/wellcomeopenres.18772.1

Lynch, M., Conery, J., Bürger, R. (1995a). Mutational meltdowns in sexual populations. Evolution, 49, 1067–1080. 10.1111/j.1558-5646.1995.tb04434.x

Lynch, M., Conery, J., Burger, R. (1995b). Mutation Accumulation and the Extinction of Small Populations. The American Naturalist, 146, 489–518. 10.1086/285812

Mackintosh, A., Laetsch, D. R., Hayward, A., Charlesworth, B., Waterfall, M., Vila, R., & Lohse, K. (2019). The determinants of genetic diversity in butterflies. Nature Communications, 10, 3466. 10.1038/s41467-019-11308-4

Mackintosh, A., Vila, R., Martin, S.H., Setter, D., Lohse, K. (2023). Do chromosome rearrangements fix by genetic drift or natural selection? Insights from *Brenthis* butterflies. Molecular Ecology, 00, 1–15. 10.1111/mec.17146

MacLachlan, R. (1893). The Decadence of British Butterflies. The Entomologist’s Monthly Magazine, 29, 132–138.

Mattila, A.L.K., Duplouy, A., Kirjokangas, M., Lehtonen, R., Rastas, P., Hanski, I. (2012). High genetic load in an old isolated butterfly population. Proceedings of the National Academy of Sciences, 109, E2496–E2505. 10.1073/pnas.1205789109

Meisner, J., Albrechtsen, A. (2018). Inferring Population Structure and Admixture Proportions in Low-Depth NGS Data. Genetics, 210, 719–731. 10.1534/genetics.118.301336

Meyermans R, Gorssen W, Buys N, Janssens S. (2020). How to study runs of homozygosity using PLINK? A guide for analyzing medium density SNP data in livestock and pet species. BMC Genomics, 29;21(1):94. 10.1186/s12864-020-6463-x.

Middleton-Welling, J., Dapporto, L., García-Barros, E., Wiemers, M., Nowicki, P., Plazio, E., Bonelli, S., Zaccagno, M., Šašić, M., Liparova, J., Schweiger, O., Harpke, A., Musche, M., Settele, J., Schmucki, R., Shreeve, T. (2020). A new comprehensive trait database of European and Maghreb butterflies, Papilionoidea. Scientific Data, 7, 351. 10.1038/s41597-020-00697-7

Mullin, V.E., Stephen, W., Arce, A.N., Nash, W., Raine, C., Notton, D.G., Whiffin, A., Blagderov, V., Gharbi, K., Hogan, J., Hunter, T., Irish, N., Jackson, S., Judd, S., Watkins, C., Haerty, W., Ollerton, J., Brace, S., Gill, R.J., Barnes, I. (2023). First large-scale quantification study of DNA preservation in insects from natural history collections using genome-wide sequencing. Methods in Ecology and Evolution, 14, 360–371. 10.1111/2041-210X.13945

Näsvall, K., Boman, J., Talla, V., Backström, N. (2023). Base Composition, Codon Usage, and Patterns of Gene Sequence Evolution in Butterflies. Genome Biology and Evolution, 15, evad150. 10.1093/gbe/evad150

Neaves, L.E., Eales, J., Whitlock, R., Hollingsworth, P.M., Burke, T., Pullin, A.S. (2015). The fitness consequences of inbreeding in natural populations and their implications for species conservation – a systematic map. Environmental Evidence, 4:5. 10.1186/s13750-015-0031-x

Neukamm, J., Peltzer, A., Nieselt, K. (2021). DamageProfiler: fast damage pattern calculation for ancient DNA. Bioinformatics, 37, 3652–3653. 10.1093/bioinformatics/btab190

Palahí I Torres, A., Höök, L., Näsvall, K., Shipilina, D., Wiklund, C., Vila, R., Pruisscher, P., Backström, N. (2023) The fine-scale recombination rate variation and associations with genomic features in a butterfly. Genome Research. 33(5), 810–823. 10.1101/gr.277414.122

Pratt, C. (1983). A modern review of the demise of *Aporia crataegi L*.: the black veined white. The entomologist’s record and journal of variation, 95, 161--166.

Preston, S.D., Liao, J.D., Toombs, T.P., Romero-Canyas, R., Speiser, J., Seifert, C.M. (2021). A case study of a conservation flagship species: the monarch butterfly. Biodiversity and Conservation, 30, 2057–2077. 10.1007/s10531-021-02183-x

Purcell, S., Neale, B., Todd-Brown, K., Thomas, L., Ferreira, M.A.R., Bender, D., Maller, J., Sklar, P., de Bakker, P.I.W., Daly, M.J., Sham, P.C. (2007). PLINK: A Tool Set for Whole-Genome Association and Population-Based Linkage Analyses. American Journal of Human Genetics, 81, 559–575. 10.1086/519795

R Core Team. (2023). R: A language and environment for statistical computing. R Foundation for Statistical Computing, Vienna, Austria.

Rhie, A., McCarthy, S.A., Fedrigo, O., Damas, J., Formenti, G., Koren, S., Uliano-Silva, M., Chow, W., Fungtammasan, A., Kim, J., Lee, C., Ko, B.J., Chaisson, M., Gedman, G.L., Cantin, L.J., Thibaud-Nissen, F., Haggerty, L., Bista, I., Smith, M., Haase, B., Mountcastle, J., Winkler, S., Paez, S., Howard, J., Vernes, S.C., Lama, T.M., Grutzner, F., Warren, W.C., Balakrishnan, C.N., Burt, D., George, J.M., Biegler, M.T., Iorns, D., Digby, A., Eason, D., Robertson, B., Edwards, T., Wilkinson, M., Turner, G., Meyer, A., Kautt, A.F., Franchini, P., Detrich, H.W., Svardal, H., Wagner, M., Naylor, G.J.P., Pippel, M., Malinsky, M., Mooney, M., Simbirsky, M., Hannigan, B.T., Pesout, T., Houck, M., Misuraca, A., Kingan, S.B., Hall, R., Kronenberg, Z., Sović, I., Dunn, C., Ning, Z., Hastie, A., Lee, J., Selvaraj, S., Green, R.E., Putnam, N.H., Gut, I., Ghurye, J., Garrison, E., Sims, Y., Collins, J., Pelan, S., Torrance, J., Tracey, A., Wood, J., Dagnew, R.E., Guan, D., London, S.E., Clayton, D.F., Mello, C.V., Friedrich, S.R., Lovell, P.V., Osipova, E., Al-Ajli, F.O., Secomandi, S., Kim, H., Theofanopoulou, C., Hiller, M., Zhou, Y., Harris, R.S., Makova, K.D., Medvedev, P., Hoffman, J., Masterson, P., Clark, K., Martin, F., Howe, Kevin, Flicek, P., Walenz, B.P., Kwak, W., Clawson, H., Diekhans, M., Nassar, L., Paten, B., Kraus, R.H.S., Crawford, A.J., Gilbert, M.T.P., Zhang, G., Venkatesh, B., Murphy, R.W., Koepfli, K.-P., Shapiro, B., Johnson, W.E., Di Palma, F., Marques-Bonet, T., Teeling, E.C., Warnow, T., Graves, J.M., Ryder, O.A., Haussler, D., O’Brien, S.J., Korlach, J., Lewin, H.A., Howe, Kerstin, Myers, E.W., Durbin, R., Phillippy, A.M., Jarvis, E.D. (2021). Towards complete and error-free genome assemblies of all vertebrate species. Nature, 592, 737–746. 10.1038/s41586-021-03451-0

Robinson, J., Kyriazis, C.C., Yuan, S.C., Lohmueller, K.E. (2023). Deleterious Variation in Natural Populations and Implications for Conservation Genetics. Annual Review of Animal Biosciences, 11, 93–114. 10.1146/annurev-animal-080522-093311

Rowe, G., Sweet, M., Beebee, T. (2017). An Introduction to Molecular Ecology, Third Edition. Oxford, Oxford University Press.

Saccheri, I., Kuussaari, M., Kankare, M., Vikman, P., Fortelius, W., Hanski, I. (1998). Inbreeding and extinction in a butterfly metapopulation. Nature, 392, 491–494. 10.1038/33136

Sarabia, C., vonHoldt, B., Larrasoaña, J.C., Uríos, V., Leonard, J.A. (2021). Pleistocene climate fluctuations drove demographic history of African golden wolves (*Canis lupaster*). Molecular Ecology, 30, 6101–6120. 10.1111/mec.15784

Solonkin, I.A., Shkurikhin, A.O., Oslina, T.S., Zakharova, E.Y. (2021). Changes in the body size of black-veined white, *Aporia crataegi* (Lepidoptera: Pieridae), recorded in a natural population in response to different spring weather conditions and at different phases of an outbreak. European Journal of Entomology, 118, 214–224. 10.14411/eje.2021.023

South, R. (1906). The Butterflies of the British Isles. London, Frederick Warne.

Teixeira, J.C., Huber, C.D. (2021). The inflated significance of neutral genetic diversity in conservation genetics. Proceedings of the National Academy of Sciences of the United States of America, 118, e2015096118. 10.1073/pnas.2015096118

Thomas, J. A. (2005). Monitoring change in the abundance and distribution of insects using butterflies and other indicator groups. Philosophical Transactions of the Royal Society B: Biological Sciences, 360, 339–357. 10.1098/rstb.2004.1585

van der Auwera, G., O’Connor, B.D. (2020). Genomics in the Cloud: Using Docker, GATK, and WDL in Terra. *O’Reilly Media*, Incorporated.

van der Valk, T., Díez-Del-Molino, D., Marques-Bonet, T., Guschanski, K., Dalén, L. (2019). Historical Genomes Reveal the Genomic Consequences of Recent Population Decline in Eastern Gorillas. Current Biology, 29, 165-170.e6. 10.1016/j.cub.2018.11.055

van Swaay, C., Wynhoff, I., Verovnik, R., Wiemers, M., López Munguira, M., Maes, D., Sasic, M., Verstrael, T., Warren, M. & Settele, J. (2010). *Aporia crataegi* (Europe assessment). The IUCN Red List of Threatened Species 2010: e.T159814A5339278. Accessed on 17 January 2024.

Wagner, D.L., Grames, E.M., Forister, M.L., Berenbaum, M.R., Stopak, D. (2021). Insect decline in the Anthropocene: Death by a thousand cuts. Proceedings of the National Academy of Sciences, 118, e2023989118. 10.1073/pnas.2023989118

Warren, M.S. (1992). The Conservation of British Butterflies, in Dennis, R.L.H. The Ecology of Butterflies in Britain. Oxford, Oxford University Press, pp. 246–274.

Warren, M.S., Maes, D., van Swaay, C.A.M., Goffart, P., Van Dyck, H., Bourn, N.A.D., Wynhoff, I., Hoare, D., Ellis, S. (2021). The decline of butterflies in Europe: Problems, significance, and possible solutions. Proceedings of the National Academy of Sciences of the United States of America, 118, e2002551117. 10.1073/pnas.2002551117

Webster, M.T., Beaurepaire, A., Neumann, P., Stolle, E. (2023). Population Genomics for Insect Conservation. Annual Review of Animal Biosciences, 11, 115–140. 10.1146/annurev-animal-122221-075025

Xue, Y., Prado-Martinez, J., Sudmant, P.H., Narasimhan, V., Ayub, Q., Szpak, M., Frandsen, P., Chen, Y., Yngvadottir, B., Cooper, D.N., de Manuel, M., Hernandez-Rodriguez, J., Lobon, I., Siegismund, H.R., Pagani, L., Quail, M.A., Hvilsom, C., Mudakikwa, A., Eichler, E.E., Cranfield, M.R., Marques-Bonet, T., Tyler-Smith, C., Scally, A. (2015). Mountain gorilla genomes reveal the impact of long-term population decline and inbreeding. Science, 348, 242–245. 10.1126/science.aaa3952

