## Supplementary Figures for "The last days of *Aporia crataegi* (L.) in Britain: evaluating genomic erosion in an extirpated butterfly"

### Supplementary Material

### The last days of *Aporia crataegi* (L.) in Britain: using museomics to evaluate genomic erosion in an extirpated butterfly

Rebecca Whitla, Korneel Hens, Casper Breuker, Timothy G. Shreeve and Saad Arif

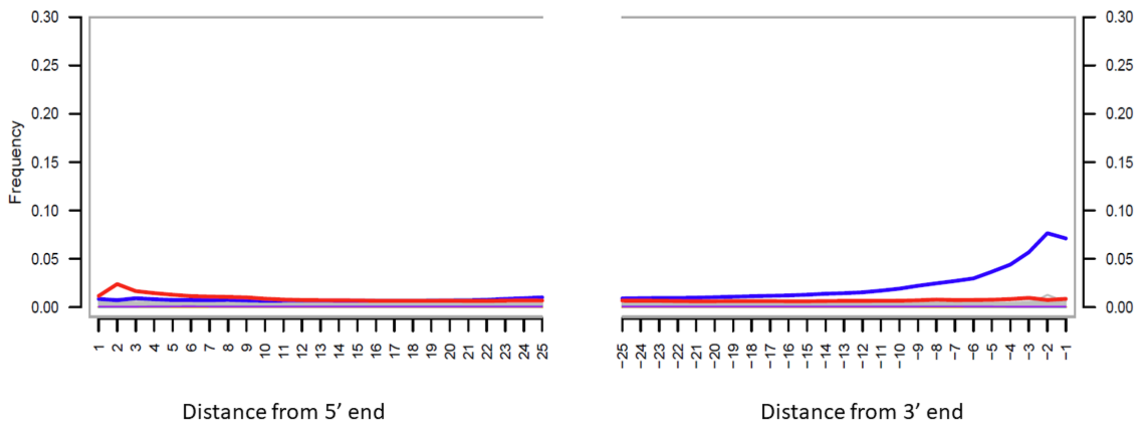


**Fig. S1** Fragment misincorporation plot of OX4 (one of the oldest specimens, collected in 1854) from DamageProfiler (Neukamm et al., 2021). Bold red lines show frequency of C to T transversions and bold blue lines show the frequency of G to A transversions along the final 25bp segments of DNA fragments.

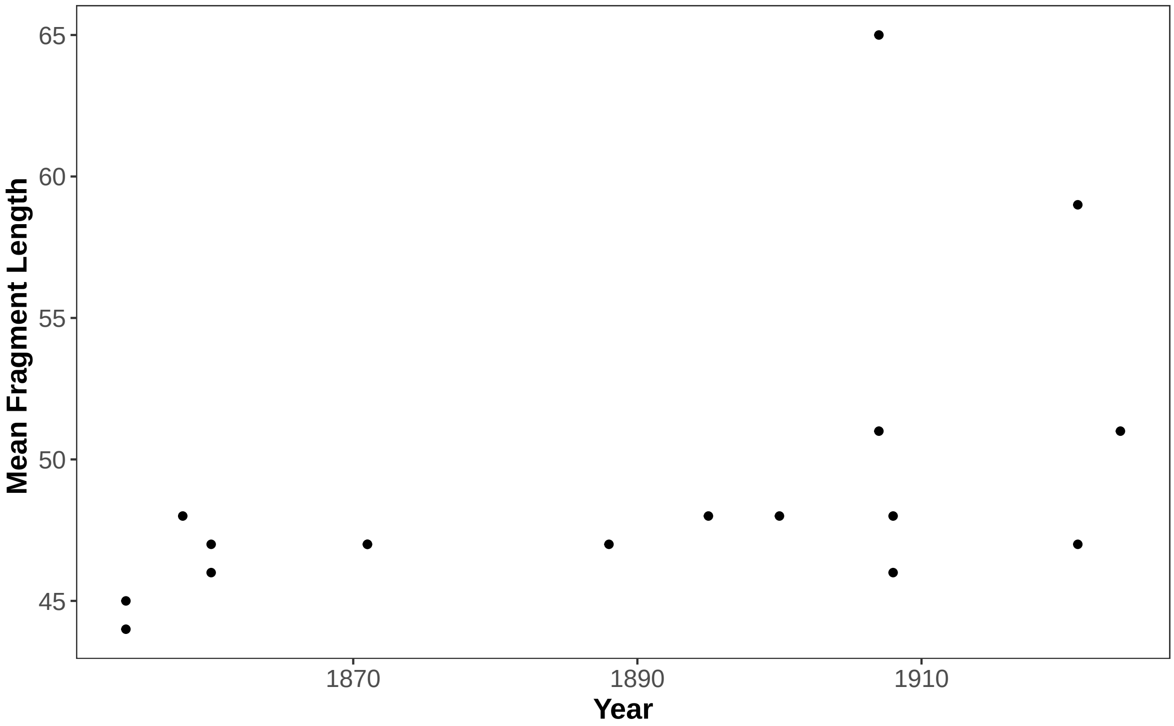


**Fig. S2** Mean fragment length by collection year. Spearmans Rank Correlation Rho = 0.607, p-value = 0.009 .

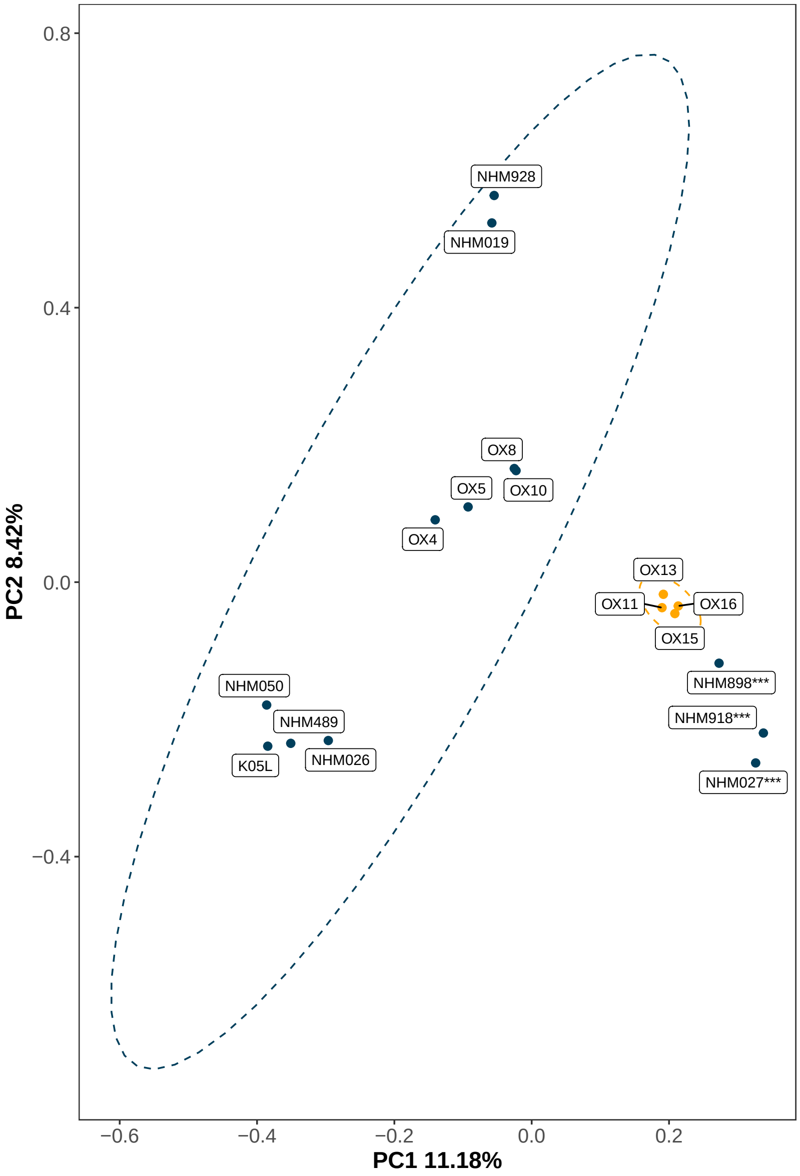


**Fig. S3** PCA of historical *A. crataegi* specimens performed using eigenvalues from genotype likelihood calls from ANGSD. Specimens NHM898, NHM918 and NHM027 group with European samples regardless of genotype calling method. This figure is related to Fig. 1.

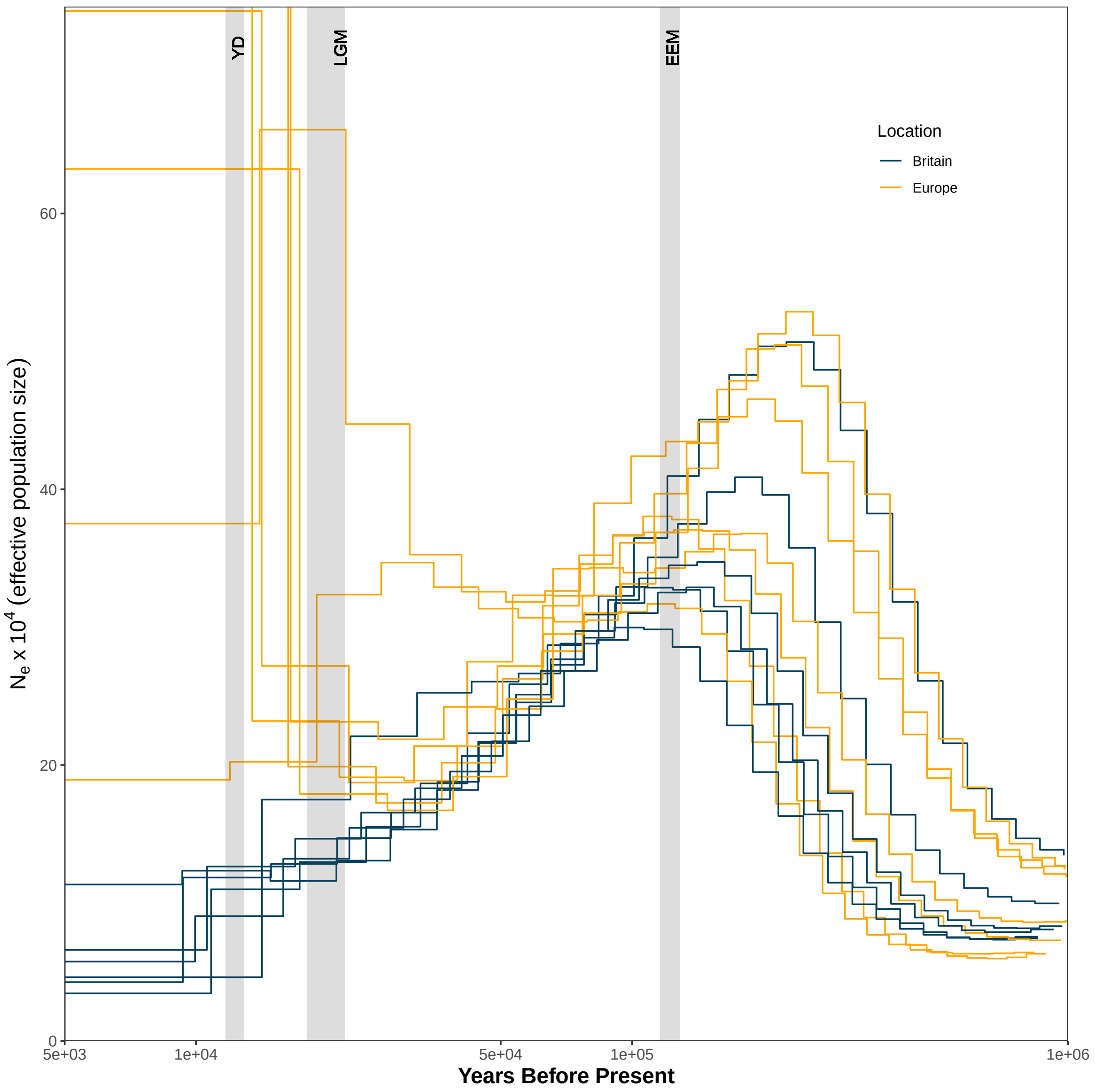


**Fig. S4** All PSMC traces for *A. crataegi* specimens. British specimens are in blue and European specimens are in yellow. All European specimens show increased Ne after the LGM in comparison to British specimens. This is related to Fig. 2.

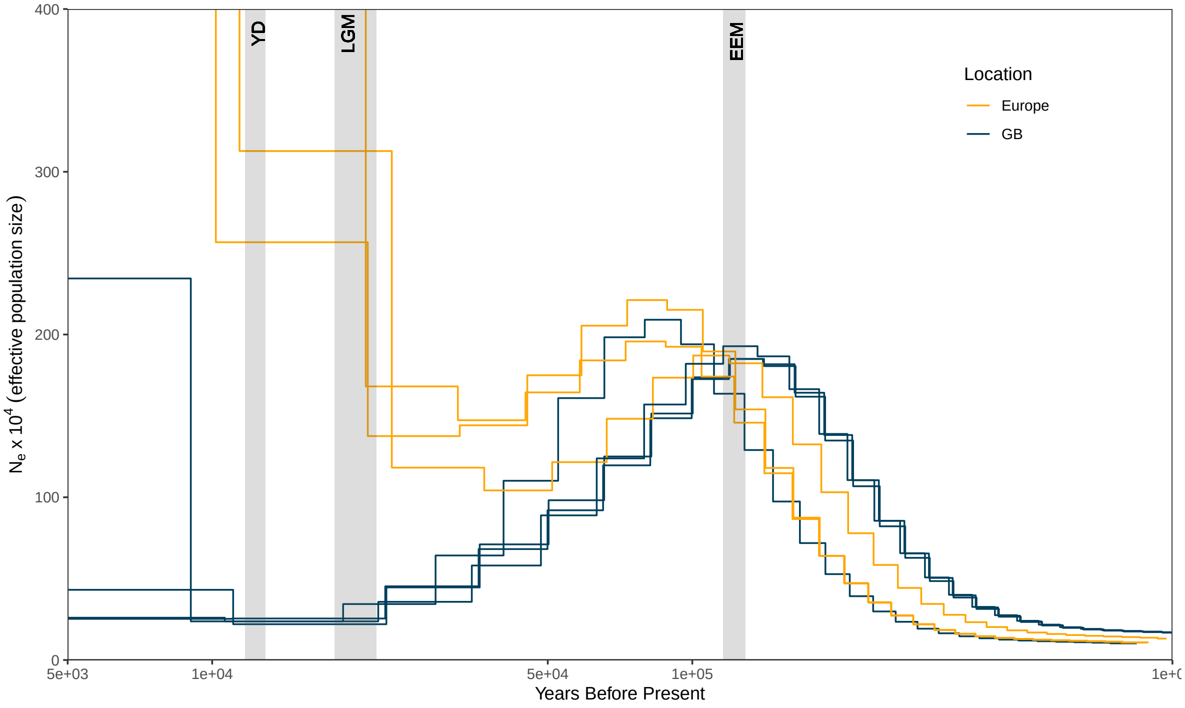


**Fig. S5** All PSMC traces for *P. icarus* specimens. British specimens are in blue and European specimens are in yellow. This is related to Fig. 2.

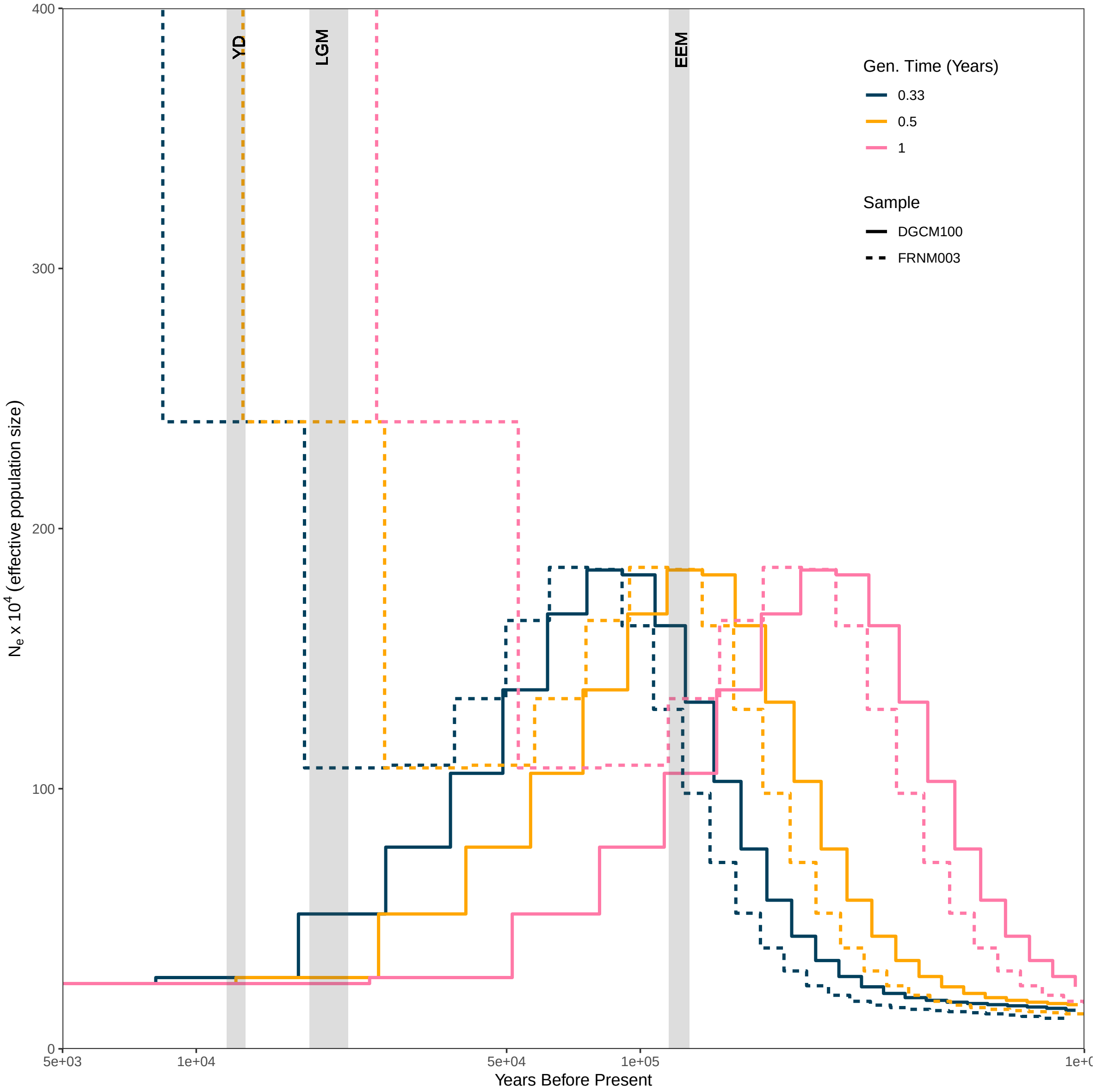


**Fig. S6** PSMC traces for *P. icarus* with generation times set to 3 broods per year (0.33), 2 broods per year (0.5) and 1 brood per year (1). This figure is related to Fig. 2.

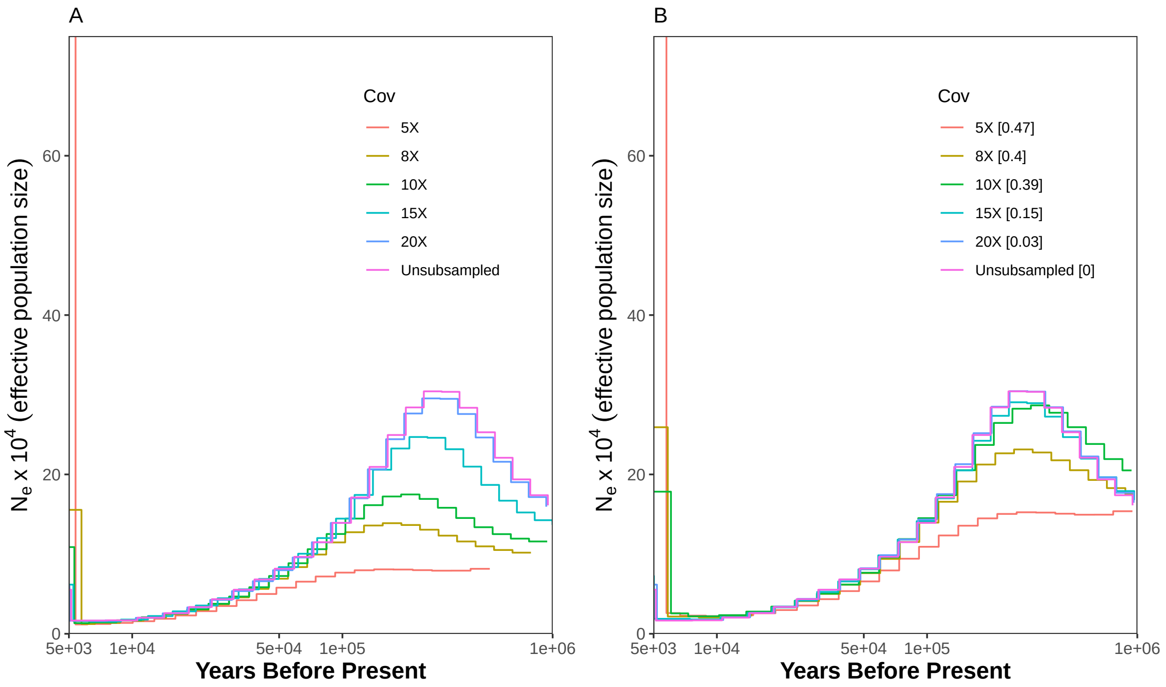


**Fig. S7 (A)**: PSMC plot showing curves using samples of differing coverages of *A. crataegi* (5X, 8X, 10X, 15X, 20X and unsubsampled). Cov=Coverage **(B)** False Negative Rate corrections used for low coverage *A. crataegi* specimens based on high coverage sample. This figure is related to Fig. 2 in main manuscript.

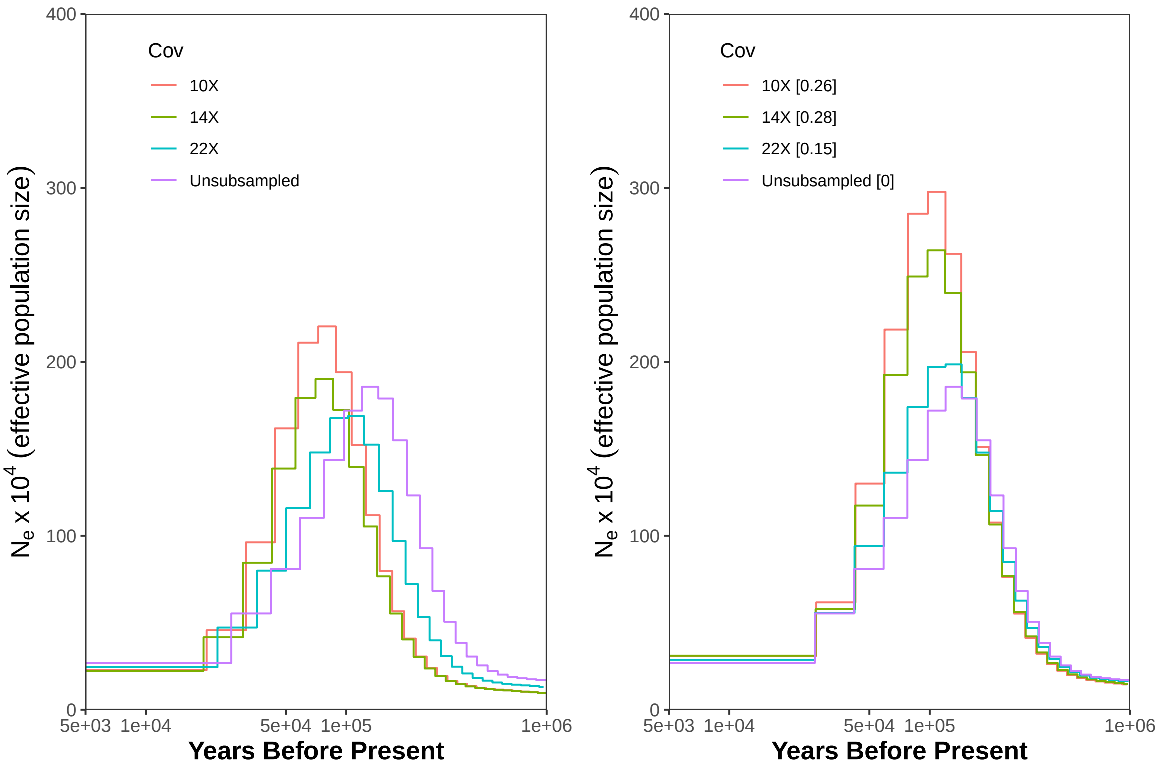


**Fig. S8 (A)**: PSMC plot showing curves using samples of differing coverages of *P. icarus* (10X, 14X, 22X and unsubsampled). **(B)** False Negative Rate corrections for lower coverage *P. icarus* specimens. This figure is related to Fig. 2.

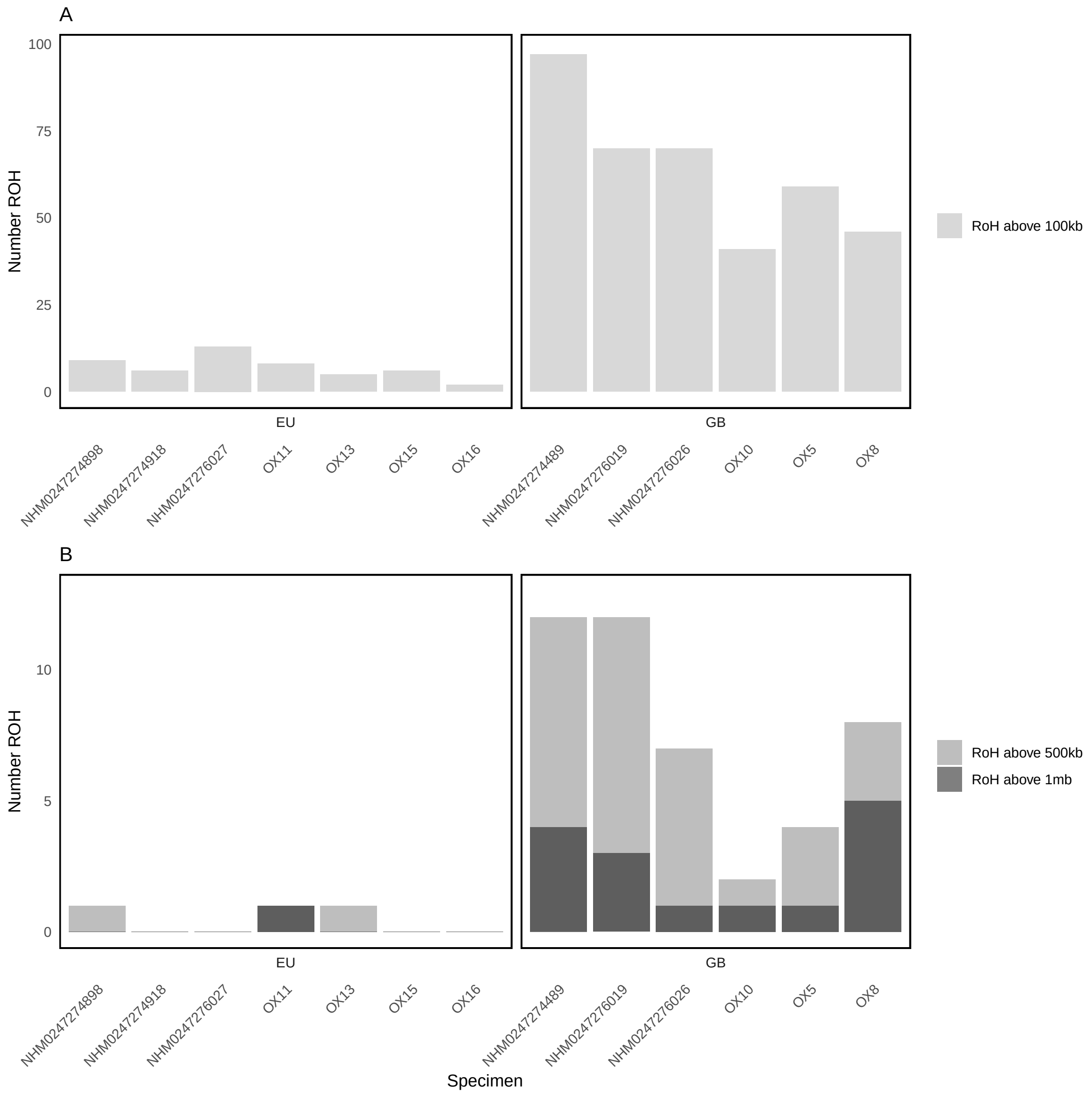


**Fig. S9 (A)**: Number of ROH above 100kb in historic *A. crataegi* samples. **(B)** Total number of ROH above 500kb and 1mb in historic *A. crataegi*.

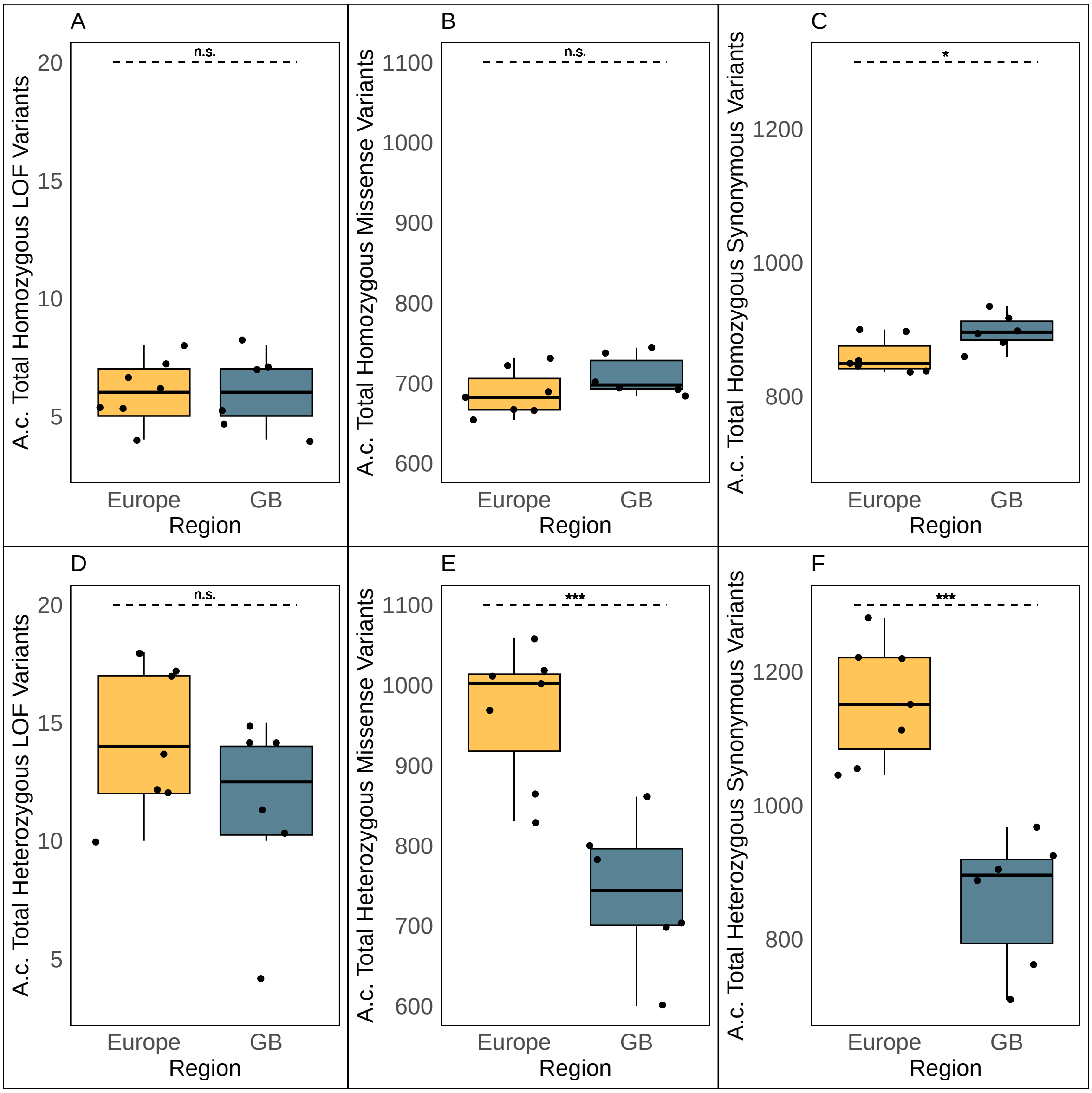


**Fig. S10** LoF, Missense and Synonymous variants in homozygous and heterozygous state in *A. crataegi*. This is related to Fig. 4.

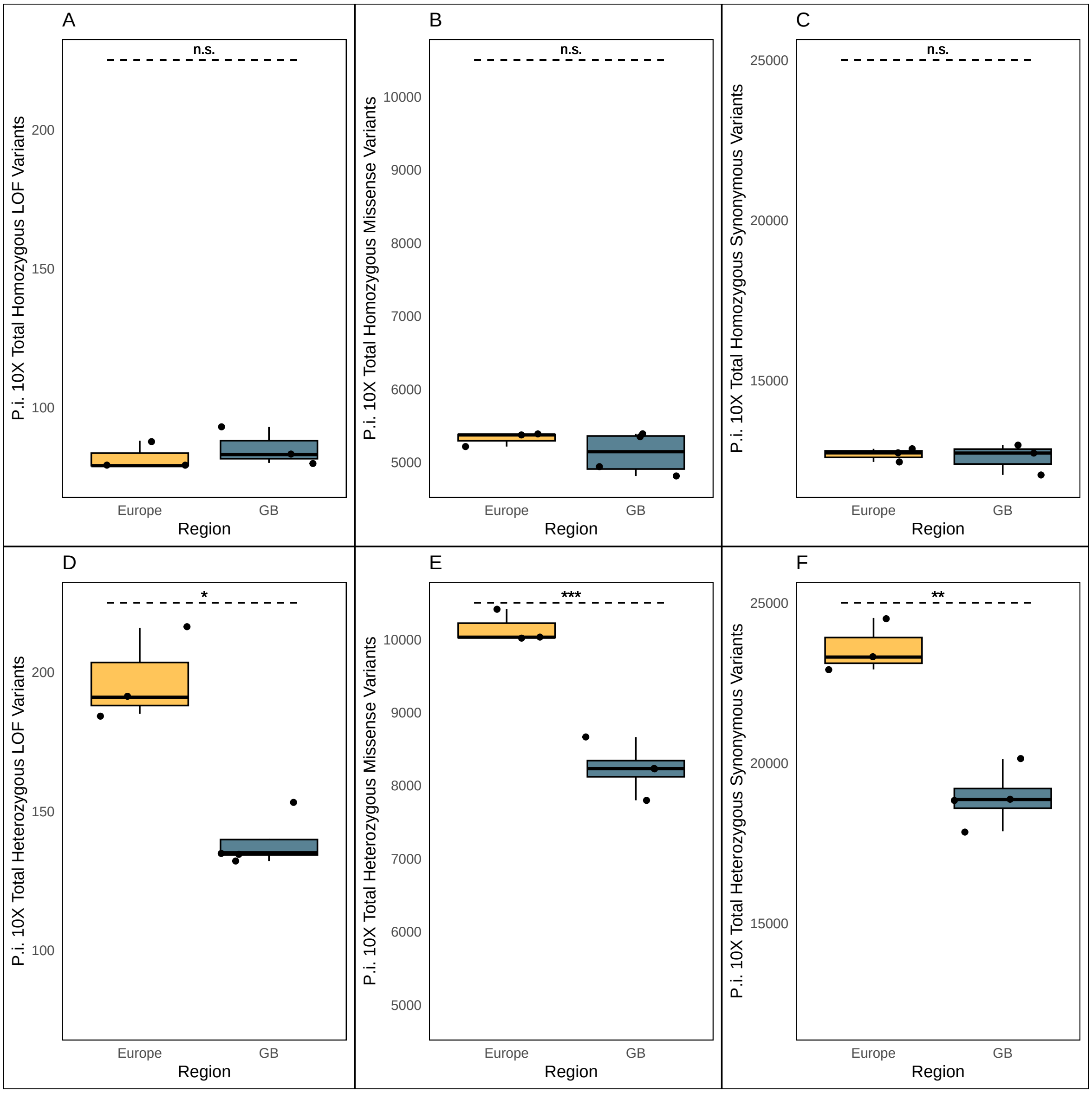


**Fig. S11** Boxplots showing total numbers homozygous and heterozygous variants in each class in European and GB *P. icarus* specimens. This is related to Fig. 4.
